# In-silico heart model phantom to validate cardiac strain imaging

**DOI:** 10.1101/2024.08.05.606672

**Authors:** Tanmay Mukherjee, Muhammad Usman, Rana Raza Mehdi, Emilio Mendiola, Jacques Ohayon, Diana Lindquist, Dipan Shah, Sakthivel Sadayappan, Roderic Pettigrew, Reza Avazmohammadi

## Abstract

The quantification of cardiac strains as structural indices of cardiac function has a growing prevalence in clinical diagnosis. However, the highly heterogeneous four-dimensional (4D) cardiac motion challenges accurate “regional” strain quantification and leads to sizable differences in the estimated strains depending on the imaging modality and post-processing algorithm, limiting the translational potential of strains as incremental biomarkers of cardiac dysfunction. There remains a crucial need for a feasible benchmark that successfully replicates complex 4D cardiac kinematics to determine the reliability of strain calculation algorithms. In this study, we propose an in-silico heart phantom derived from finite element (FE) simulations to validate the quantification of 4D regional strains. First, as a proof-of-concept exercise, we created synthetic magnetic resonance (MR) images for a hollow thick-walled cylinder under pure torsion with an exact solution and demonstrated that “ground-truth” values can be recovered for the twist angle, which is also a key kinematic index in the heart. Next, we used mouse-specific FE simulations of cardiac kinematics to synthesize dynamic MR images by sampling various sectional planes of the left ventricle (LV). Strains were calculated using our recently developed non-rigid image registration (NRIR) framework in both problems. Moreover, we studied the effects of image quality on distorting regional strain calculations by conducting in-silico experiments for various LV configurations. Our studies offer a rigorous and feasible tool to standardize regional strain calculations to improve their clinical impact as incremental biomarkers.

## 1 Introduction

Cine imaging of the left ventricle (LV) is a vital component in the diagnostic protocol for numerous cardiac diseases [1, 2]. Organ-level biomarkers, such as the ejection fraction (EF), end-diastolic volume (EDV), and stroke volume (SV), are key determinants of ventricular dysfunction [2, 3]. Despite the diagnostic importance of these traditional markers at advanced stages of cardiac disease, they might not be sufficiently sensitive to the progression of many diseases at their early-stage development [4, 5]. In contrast, cardiac strains measured from cine images are found to be more sensitive indicators of cardiac dysfunction at an early stage [6, 7]. Strain components, commonly expressed along the anatomical axes, i.e., the radial, circumferential, and longitudinal (RCL), have shown improved sensitivity to multiple cardiovascular diseases [8, 9, 10]. Additionally, principal strain analysis has provided insight into architectural alterations in the LV [9], thus offering the potential to delineate the LV structure-function relationship. Nonetheless, the strong heterogeneous variations in cardiac deformation, known as strain regionality, and significant out-of-plane motion of the LV present challenges. Subsequently, the utility of global strain markers, including GLS, is critically affected by the ability to accurately estimate the complete spatial and temporal variations in cardiac motion, referred to as four-dimensional (4D) quantification of the strains. Although the advancement of cardiac imaging and the capability of multi-plane acquisitions have indeed enabled the high-fidelity characterization of regional strains [11], the absence of ground-truth measurements of 4D cardiac motion leaves existing tools of strain analyses insufficiently validated.

Cardiac strains, for the most part, are estimated via ultrasound and cardiac magnetic resonance (CMR) imaging [12, 13]. A common benchmark in estimating regional strains is the application of tagged CMR images to track cardiac deformation using displacement encoding (DENSE) [14] and harmonic phasing [15] algorithms. The complex protocols involved in tag deposition and tracking may pose challenges for feasible clinical translation, especially when 4D strain calculations are of interest. Motion-tracking approaches such as speckle tracking echocardiography (STE) [13] and feature tracking (FT) [16] offer potential clinical utility with lower startup costs, making them promising additions to strain calculation methodologies. STE using brightness-mode (B-mode) ultrasound imaging is perhaps the most prevalent application of regional strain calculations. Additionally, FT algorithms have offered higher fidelity by estimating three-dimensional (3D) cardiac strains using cine CMR images. Despite potential benefits, clinical adoption of regional strains remains limited, as technological factors, including differences in vendors, operators, and post-processing schemes, impact consistency between studies, leading to substantial calculation variabilities [16]. For instance, Verzhbinsky et al. [17] discuss the effects of image quality as a potential source of variation between studies and demonstrate the effect of suboptimal image resolution on the underestimation of radial strains. Indeed, imaging aspects such as spatial and temporal resolution, signal-to-noise ratio (SNR), and artifacts contribute to spurious strain estimation. For example, cine CMR images are susceptible to poor spatial resolution, challenging the delineation of cardiac motion along its long axis [16, 18]. Thus, these variabilities can significantly hinder the reliable and consistent application of regional strain calculations in clinical practice. We hypothesize that the reliability of current strain calculation tools can be substantially improved through our proposed in-silico validation protocols.

Numerous studies have underscored the benefits of establishing a validation benchmark in improving the reliability of regional strain calculations [11, 17, 19, 20]. Perhaps the most commonly implemented protocol is using phantoms to validate imaging and post-processing methods [11]. Traditional approaches have been restricted to validating strain calculations under basic kinematic movements, such as rotation [21, 22] and inflation [23]. Recent efforts in cardiac motion validation have shown the utility of *in-vitro* phantoms in evaluating myocardial mechanics under more complex loading conditions. In their study, Wang et al. [24] introduced a novel kinematic analysis framework to compute 3D DENSE-derived strains, which were subsequently subjected to validation using a cylindrical gel phantom subjected to axial compression. Overall, the regional variations in the myofiber organization [25, 26], tissue compressibility [27, 28] and torsional mechanics [21] challenge the accuracy of motion tracking [11, 24] and necessitate the development of a validation protocol that could recapitulate all the regional high-fidelity features. Along these lines, Kolawole et al. [29] designed an anatomically detailed gel phantom to closely resemble the passive mechanical and geometric properties of the heart, with cardiac motion driven through an external pump.

The material parameters were established using a ground-truth mechanical characterization of the gel stiffness, and the diastolic filling was mimicked via the injection of a fluid volume. Despite advancements in enhancing the overall fidelity of phantom-based validation, in-vitro validation encounters three major limitations: (a) inadequate representation of accurate cardiac motion, exacerbated by oversimplified LV kinematics and lack of myofiber network, (b) impractical integration of subject-specific pressure-volume (P-V) readings with the phantom, and (c) challenges in quantification of ground-truth deformation in case of non-idealized geometry and motion of in-vitro phantoms. In contrast, *in-silico* finite element (FE) simulations offer to incorporate subject-specific geometries, myocardial architecture, and pressure distributions, holding the potential to establish a robust kinematic benchmark [30, 31, 32, 33, 34] for validating cardiac strains. These FE models have demonstrated a high level of accuracy in capturing myocardial mechanics in both healthy and pathological conditions [25, 26, 35]. Thus, the utilization of in-silico phantoms derived from biomechanical models of the heart shows promise in addressing the limitations of the current in-vitro protocols. However, the implementation of FE-based tools for validating regional strain calculations remains critically understudied.

In this study, an in-silico phantom to validate regional cardiac strains by generating synthetic images derived from FE simulations is proposed (Fig. 1). Since cardiac motion tracking is often performed across several short-axis (SA) slices of the LV, phantom images were sampled from different sectional planes of the deforming geometry. Phantoms were synthesized from the geometry at multiple time frames by prescribing FE-derived displacements over the cardiac cycle. An in-house non-rigid image registration (NRIR) framework was devised to calculate four-dimensional (4D) cardiac strains., i.e., the complete spatiotemporal deformation of the LV at end-systole (ES) relative to end-diastole (ED). Validation was performed by comparing strains derived from FE (considered to be “ground truth”) and those from NRIR at ES in the RCL axes. Further, we conducted sensitivity analyses using phantoms of various configurations to investigate the contributors to spurious imaging-derived regional strain estimation. To the best of our knowledge, this is the first implementation of an FE-derived kinematic benchmark of cardiac motion. The versatility of the FE-derived phantoms in representing the complexities of myocardial mechanics shows high potential in standardizing regional strain calculations for inclusion as routine clinical cardiac function assessment.

**Fig. 1.**
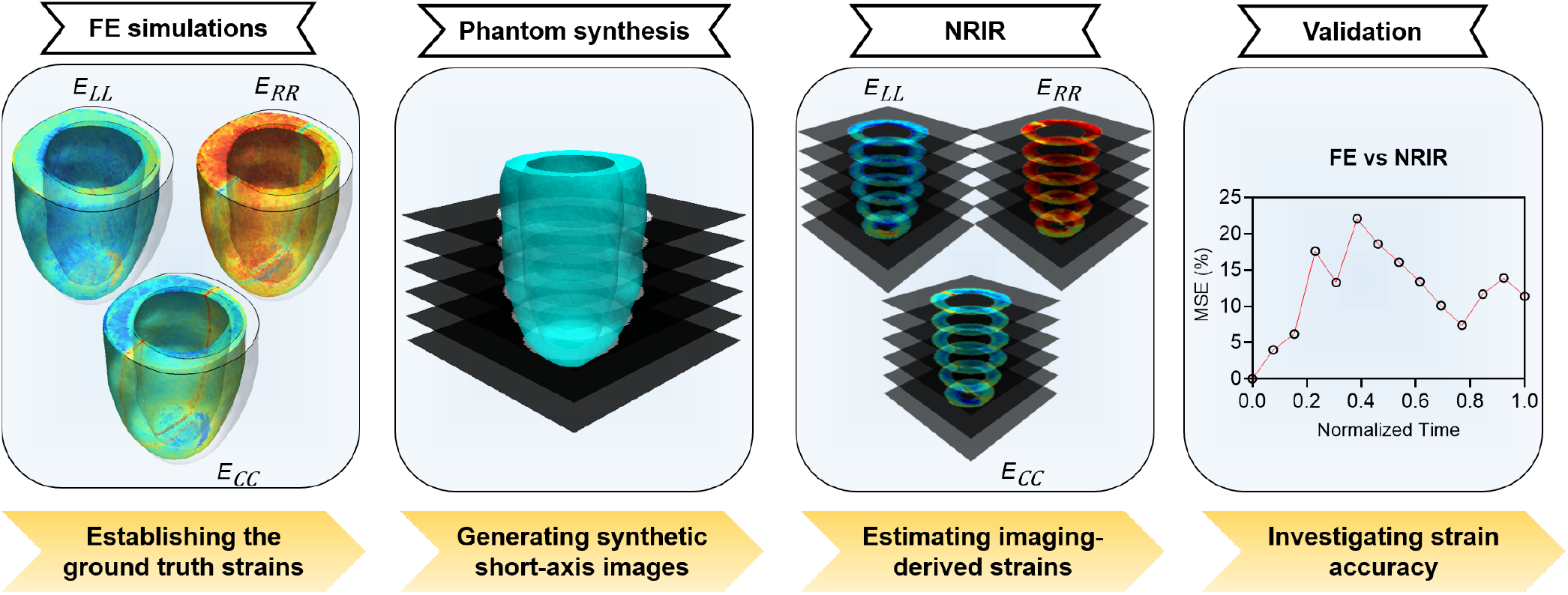
Schematic outlining the stages in the validation of cardiac strain imaging. (1) Finite element (FE) simulations of left ventricular kinematics, subsequent (2) generation of in-silico phantoms, (3) non-rigid image registration (NRIR) solutions of radial (*E*_*RR*_), circumferential (*E*_*CC*_) and longitudinal (*E*_*LL*_) strains, and (4) validation analysis.

## 2 Methods

Torsion is a prominent component of LV motion, playing a key role in the physiological function of the heart. Capturing the LV torsion kinematics accurately must be central to developing cardiac strain calculation methods. Accordingly, we first examined our approach for an idealized geometry of a thick-walled hollow cylinder under pure torsion where an exact solution for the strain is obtainable (Fig. 2A; Section 2.1). The FE simulations of the hollow cylinder under torsion were used to extract strains and verified against the analytical solution (Appendix A1). Next, an image-based FE model of a murine heart with subject-specific material and loading parameters was used to design a heart model phantom. As murine hearts exhibit rapid cardiac motion due to underlying fast cardiac metabolism, further challenging 4D strain calculations compared to the human heart, we developed the in-silico phantom for a murine heart to establish a rigorous validation protocol for 4D strain calculation algorithms. The myocardium was modeled as an incompressible hyperelastic transversely isotropic solid with a mouse-specific fiber architecture. The LV was subjected to loading via an organ-level P-V loop to recapitulate the in-vivo cardiac motion (Figs. 2B-D; Section 2.2). In addition to the murine heart model phantom, a human-specific heart model phantom was created to present the potential clinical feasibility of the proposed phantom (Appendix A2).

**Fig. 2.**
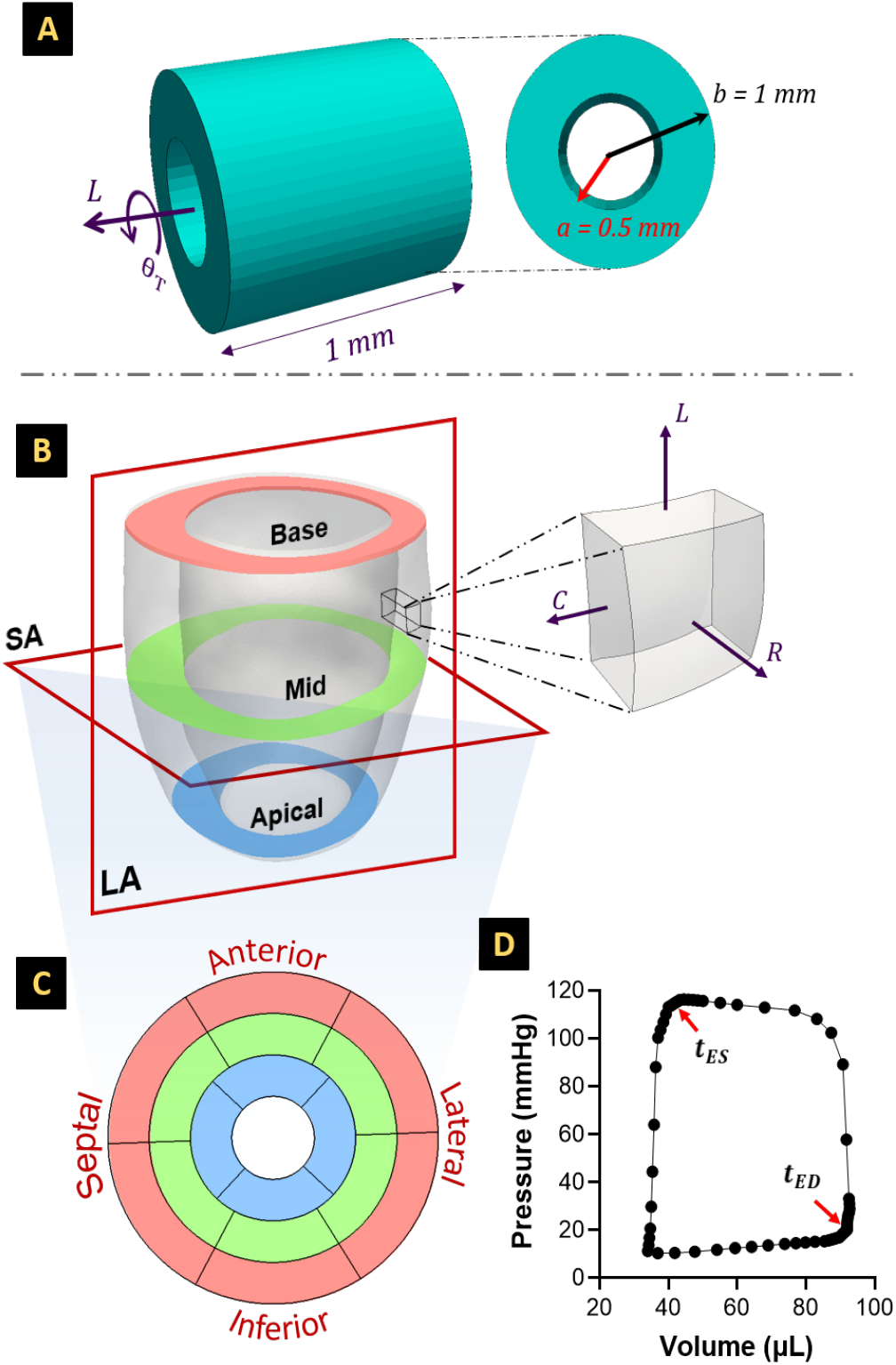
Finite element (FE) set up for simulations of (A) a thick hollow cylinder of inner and outer radius *a* = 0.5 mm and *b* = 1 mm, respectively, and height = 1 mm subjected to pure torsion via axial rotation about the longitudinal axis. The geometry was rotated over fifty equal load increments up to a maximum twist angle (*θ*_*T*_ = 30°). (B-D) A murine left ventricle (LV) under subject-specific kinematic loading. (B) Geometric reconstruction of the LV using subject-specific cardiac magnetic resonance (CMR) images, with a representative cube element of the cardiac tissue highlighting the radial, circumferential, and longitudinal axes. (C) Projection of the base, mid, and apical slices of the LV onto a two-dimensional AHA segmentation map, with the outermost and innermost circles corresponding to the LV base and apex, respectively. (D) The geometry was loaded via a mouse-specific pressure-volume loop for sixty load increments. *t*_*ED*_: end-diastolic time frame, *t*_*ES*_: end-systolic time frame.

### 2.1 Pure torsion of a hollow cylinder

FE simulations were conducted to determine the deformation of a thick hollow cylinder under pure torsion. A hollow cylinder of inner and outer radius *a* = 0.5 mm and *b* = 1 mm, respectively, and height = 1 mm was created and meshed using linear tetrahedral elements. An adaptive mesh strategy was implemented, wherein the element length was fixed at 0.1 mm in the inner and outer edges of the cylinder. The cylinder was modeled as an incompressible hyperelastic material following the Mooney-Rivlin invariant-based formulation. The strain energy function (*ψ*) characterizing the cylinder behavior was given by:

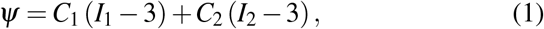

where *I*_1_ and *I*_2_ are the first and second invariants of the right Green deformation tensor. The material parameters, *C*_1_ and *C*_2_, were set to be 0.2 MPa and 0.005 MPa, respectively. The material parameters represent a hypothetical rubber-like cylinder to generate high resistance to compression through *C*_1_ while modulating the shear response through *C*_2_. Torsional loading was imposed by applying a kinematic rotation to the top of the cylinder about the z-axis while constraining the deformation of the base in all directions. The maximum twist angle was restricted to *θ*_*T*_ =30°, and the cylinder was rotated incrementally over fifty equal increments. Geometry construction, meshing, and simulations were performed in ABAQUS (Dassault Systemes Simulia Corporation, Providence, RI.).

### 2.2 Mouse-specific LV kinematics

LV kinematics was generated through forward FE simulations of a mouse-specific model of the heart to obtain cardiac displacements and strains. The LV geometry was reconstructed using CMR images, and FE simulations for a full cardiac cycle were conducted after assigning material behavior and prescribing boundary conditions. The details of similar rodent-specific in-silico models have been described extensively in our previous studies [25, 36], and are summarized below.

#### 2.2.1 Geometry reconstruction

The geometry of the LV was reconstructed at ED using eight short-axis (SA) CMR scans of a murine model. The cardiac wall was isolated by segmenting the endo- and epicardium at each SA slice using 2D contours. Segmentation was performed in Segment version 3.0 R8531 [37]. The pixels corresponding to the endo- and epicardium were connected using the three-dimensional Delaunay triangulation algorithm, and the resulting surface geometries were constructed using the Python interpretation of the Visualization Toolkit (VTK) [38]. Subsequently, the resulting surface was converted into a volume and discretized into linear tetrahedral elements. Discretization was performed as follows: the space between the two boundaries, i.e., endo- and epicardium, was filled through an adaptive mesh strategy, with the minimum element size maintained at 0.5 mm. The distance between nodes at the boundaries was specified at 1 mm, measured as the length of the hypotenuse of the linear tetrahedral element. Discretization and mesh refinement were performed in Materialize Mimics Innovation Suite (Materialise, Leuven, Belgium).

#### 2.2.2 Material model

Following reconstruction, myofiber architecture was assigned to the LV using a Laplace-Dirichlet rule-based algorithm [25]. A transversely isotropic hyperelastic material model, characterized by a single fiber direction, was assigned to the LV geometry [26]. The myocardial wall was divided into four layers, with the helicity of the fibers *θ* assumed to vary linearly from negative orientation at the epicardium (*θ*_*epi*_ = −42°) to positive orientation at the endocardium (*θ*_*endo*_ = 84°) The preferred myofiber angle was determined from our prior work in ex-vivo histological studies in wild-type mice [39]. Active and passive stress responses within the myocardium were individually described by:

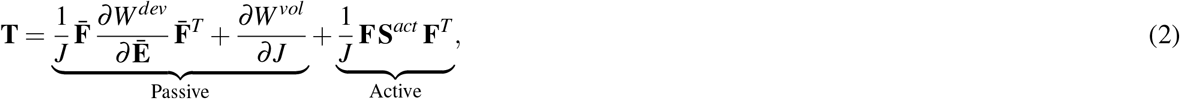

where **T** is the Cauchy stress tensor, **F** is the deformation gradient, *J* denotes the deformation volumetric changes, and deviatoric part of **F** is represented by 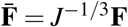. The passive stress was quantified using a Fung-type strain energy function as:

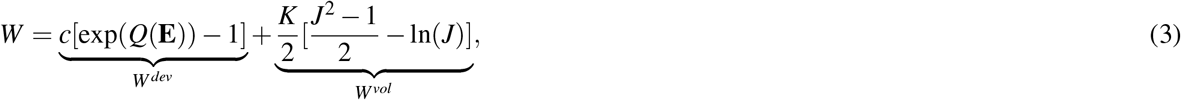

where **E** is the Lagrangian formulation of the Cauchy-Green strain tensor, *W*^*dev*^ and *W*^*vol*^ represent the deviatoric and volumetric components of *W*, respectively, *K* is the bulk modulus, and *Q* is a quadratic form of an incompressible hyperelastic material expressed as:

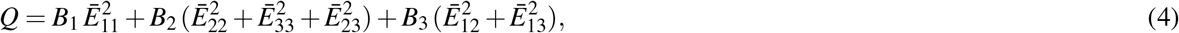

where the subscripts denote the local preferred material directions in Cartesian coordinate system {**e**_1_, **e**_2_, **e**_3_}, with the myofiber direction being along **e**_1_. In Eqs. 3 and 4, *c* is a positive stiffness parameter, *B*_1_, *B*_2_, and *B*_3_ are dimensionless constants characterizing the local anisotropy in the myocardium, and *K* is the bulk modulus. All passive properties were estimated using the average stress-strain response (n = 6) from biaxial testing of healthy mouse myocardium ex vivo [39]. The active stresses in the Eq. 2 were estimated using the second Piola–Kirchhoff active stress tensor **S**^*act*^ as:

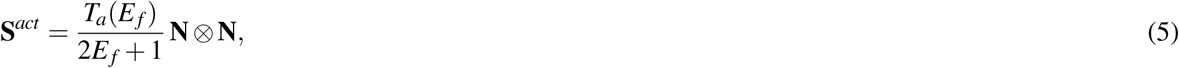

where *T*_*a*_(*E*_*f*_) is a stress-like positive function of the strain in the fiber direction **N** given by *E*_*f*_ = **N** · **EN**. We chose the following form for *T*_*a*_(*E*_*f*_)

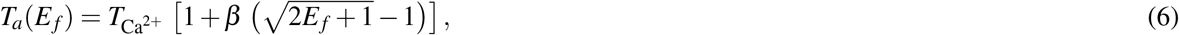

where the calcium-activated force 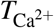 developed within the resting myofibers [40] increases by a positive factor *β* when the myofibers are stretched to the strain *E*_*f*_, obeying the Frank-Starling relationship. The experimental measurements and inverse problem approach were used to determine the passive (Eqs. 3, 4) and active parameters (Eq. 6) as described by Mendiola et al. [36].

### 2.3 Phantom synthesis

Phantom images were synthesized *in silico* from the 3D FE-derived displacements and consisted of the following steps for both torsion and LV kinematic problems (Supplementary Fig. S1 and Fig. 3, respectively). The connectivity data of the meshed geometry was initially used to create an unstructured VTK grid in the undeformed state (**X** ∈ ℝ^3^(*x, y, z*)). Grid deformation was simulated by applying the FE-derived Cartesian displacement vectors onto each node at every load increment, resulting in individual grids for each load increment. These grids were sectioned at incremental positions along the z-axis and projected onto a uniform grid of predefined parameters, thus creating a rasterized image representation for the 3D grid. Subsequently, synthetic 2D images along specific sectional planes were formulated as a linear system,

**Fig. 3.**
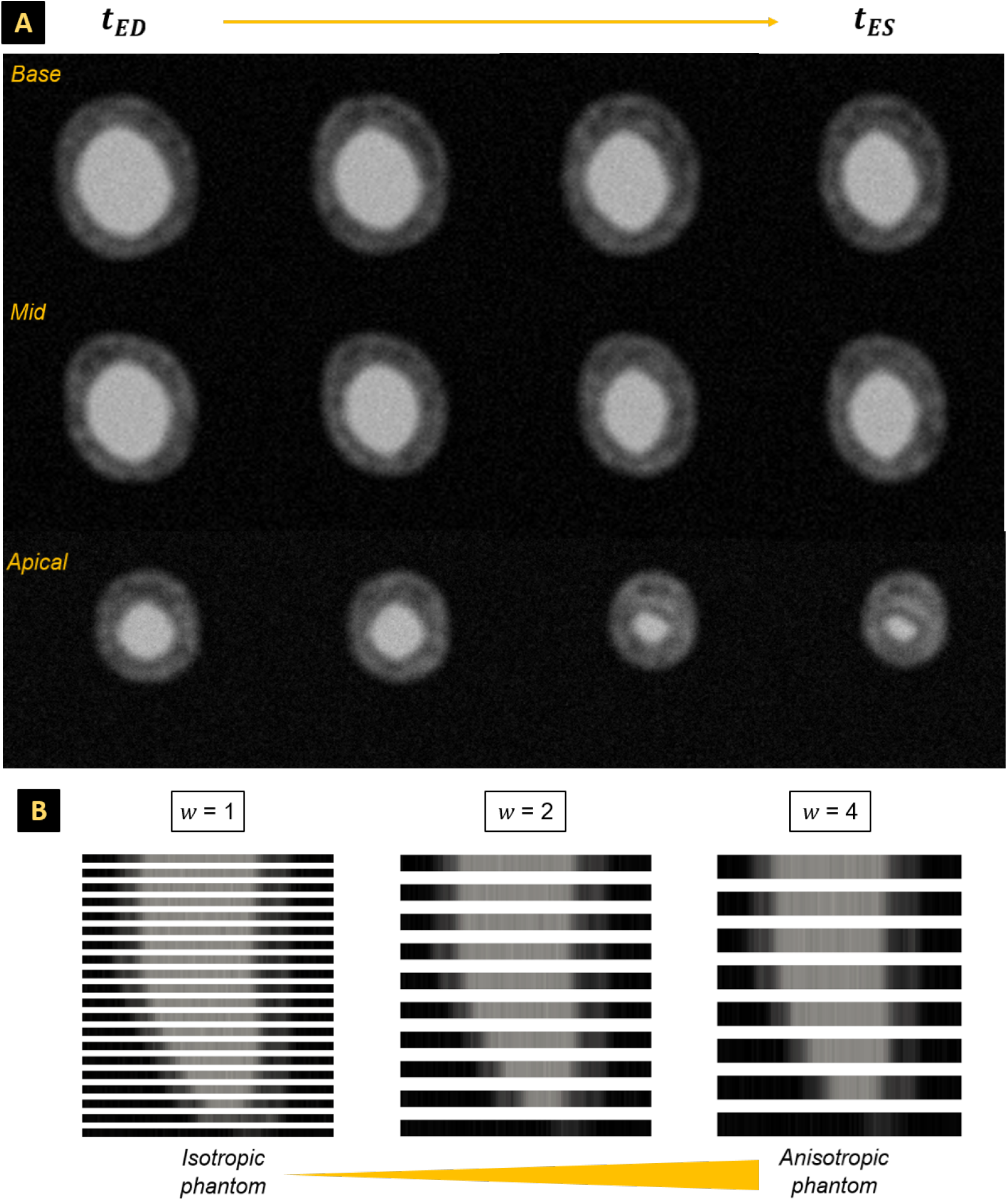
(A) Grayscale images of the in-silico heart model phantom at various short-axis sectional planes of the left ventricle created using FE-derived Cartesian displacement vectors at several time frames between end-diastole (ED) and end-systole (ES). These images correspond to a phantom of spatial resolution = 1*×*1 mm^2^ and slice thickness (*w*) = 1 mm. (B) The through-plane reconstruction of the long-axis of the left ventricle using a stack of short-axis slices at end-diastole. Three slice thicknesses were chosen: *w* = 1, 2, and 4 mm, such that (left to right) the through-plane reconstruction ranged from an isotropic phantom (*w* = 1 mm) to a strongly anisotropic phantom wherein the through-plane resolution is four times the in-plane resolution (*w* = 4 mm). Additionally, the reduction in the field of view (i.e., right to left) was compensated by increasing the number of SA slices in the phantoms. *t*_*ED*_: end-diastolic time frame, *t*_*ES*_: end-systolic time frame.

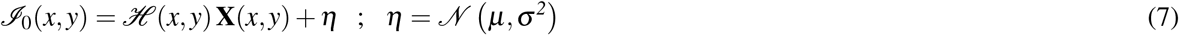

where ℐ_0_ is obtained via the convolution of **X**(*x, y*) with a function, ℋ (*x, y*) and by adding noise, *η. η* was modeled as a normal distribution 𝒩 of mean *µ* and standard deviation *σ*, respectively. Since the beam formation of most imaging systems can be modeled as sinusoidal waves, ℋ (*x, y*) was formulated as

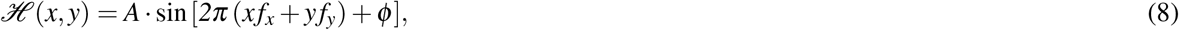

where *A* is the amplitude of the wave, ϕ is the phase, and *f*_*x*_ and *f*_*y*_ are the wave numbers in the *x* and *y* directions, respectively. The mean gray level of the cardiac muscle was considered to be 30% lower than the blood pool. As a result, a stack of short-axis images describing the deformation of the geometry was obtained. Given the predetermined number of load increments used in the FE solutions, each stack was conceptualized as an individual temporal frame within an imaging sequence (Fig. 3A). Finally, the phantoms were exported as .tiff images using the lossless Lempel-Ziv-Welch compression algorithm with grid information (plane position, plane normal, projection angles, pixel spacing, resolution, etc.) passed as metadata.

#### 2.3.1 Synthetic image generation

Multiple phantoms were created by varying imaging parameters such as resolution and SNR. The spatial resolution of the phantoms was controlled by varying the spacing of the uniform grid (in-plane resolution) and slice thickness (through-plane resolution), and the temporal resolution was manipulated by skipping load increments. To increase the field of view of each phantom, the reduction in slice thickness was compensated by an increase in the number of planes describing the geometry, i.e., a reduction in the slice gap. Thus, the spatial resolution was varied such that the phantom configurations ranged from strongly anisotropic to completely isotropic. Here, isotropy refers to equality in the through- and in-plane resolutions. Additionally, the SNR of the phantom images was varied by adding noise intensity, modeled as a normal distribution. The noise intensity was controlled by varying the standard deviation of the normal distribution (*σ*) in the image formation process. Phantoms of various configurations were created for the torsion and LV kinematic problems and are summarized in Tables 1 and 2, respectively. The field of view describing the LV LA in the phantoms of lower slice thickness (*w* = 1 mm and *w* = 2 mm) was improved by increasing the number of SA slices describing the LV (Fig. 3B). Additionally, a phantom more representative of a murine CMR phantom was created by setting the through-plane resolution to four times the in-plane resolution (*w* = 4 mm; Fig. 3B).

**Table 1.**
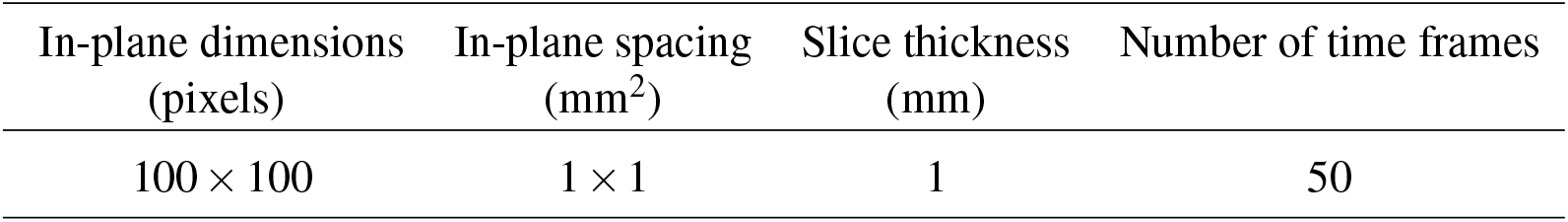
Summary of the image parameters used to create phantoms for the problem describing pure torsion of a thick-walled hollow cylinder (section 2.1). Various configurations of the phantom images were created through different combinations of pixel spacing, slice thickness, and temporal resolution (number of time frames).

**Table 2.** Summary of the image parameters used to create phantoms for the problem describing mouse-specific LV kinematics (section 2.2). Various configurations of the phantom images were created through different combinations of pixel spacing, slice thickness, temporal resolution (number of time frames), and noise intensity (varying the standard deviation of a normal distribution), *σ*: standard deviation (SD) of a normal distribution.

| In-plane dimensions<br>(pixels) | In-plane spacing<br>(mm<sup>2</sup>) | Slice thickness<br>(mm) | Number of time frames | Noise |
| --- | --- | --- | --- | --- |
| 100 × 100 | 2 × 2 | 1,2,4 | 15,30,60 | $\sigma = 1, 2, 3$ |
| 100 × 100 | 1 × 1 | 1,2,4 | 15,30,60 | $\sigma = 1, 2, 3$ |
| 100 × 100 | 0.5 × 0.5 | 1,2,4 | 15,30,60 | $\sigma = 1, 2, 3$ |

#### 2.3.2 Image quality analysis

The spatial frequency response (SFR) for an arbitrary LA plane was calculated for each LV phantom by evaluating the 2D Fourier transform of the line spread function of each image (Fig. 3B). The horizontal and vertical spatial frequencies were normalized, and a normal distribution curve was fitted to quantitatively investigate the spatial resolution. Image sharpness was also evaluated by convolving each image with its corresponding SFR.

### 2.4 Image registration

In-silico experiments were conducted to derive deformation from the phantom images. Non-rigid image registration (NRIR) using the diffeomorphic demons algorithm was employed to quantify deformation at the maximum deformed state for problems 2.1 and 2.2 [41]. The deformation for problem 2.2 was estimated at the time frame corresponding to ES with the end-diastolic frame as a reference. The optimization-based image registration problem was implemented to find the best transformation in aligning a reference frame, ℱ and a moving image, ℳ through the minimization of the cost function, 𝒟 as:

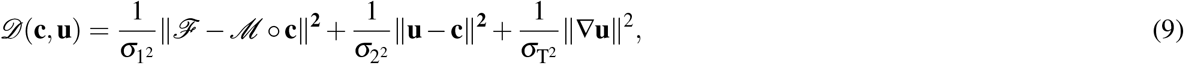

where *σ*_1_ and *σ*_2_ are the noise intensity parameter and spatial uncertainty parameter, respectively, and *σ*_T_ is a regularization factor. Here, the parametric transformation (**u**) refers to the Cartesian displacement vector at each pixel between two consecutive time frames, and **c** denotes the corresponding non-parametric spatial transformation.

### 2.5 Strain analysis

Following in-silico simulations, the large deformation formulation was used to derive a finite-strain strain measure for both the FE- and image-derived Cartesian displacements in both the torsion and LV kinematic problems (Sections 2.1 and 2.2). As finite-strain problems, different measures of strains (i.e., Eulerian or Lagrangian) can be calculated for both the FE- and image-served displacements. Both Eulerian and Lagrangian measures were used to demonstrate the generality of the chosen strain measure. Strains in the torsion problem (Section 2.1) were calculated using the Eulerian approach as:

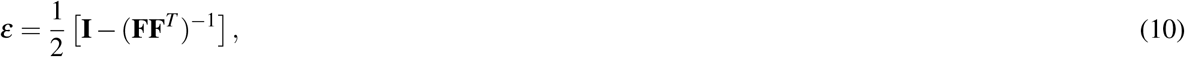

where *ε* is the Eulerian strain tensor, **F** is the total deformation gradient, and **I** is the identity matrix. For the mouse-specific LV kinematics (Section 2.2), the common Lagrangian strain measure (**E**) was calculated as:

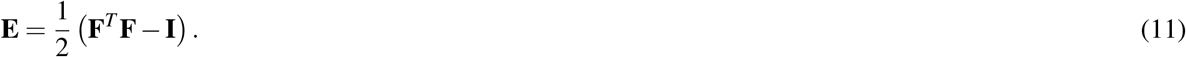

For both problems, the derivation of **F** to calculate image-derived strains was performed through the propagation of the deformation gradient between consecutive images (**F**_i_) such that

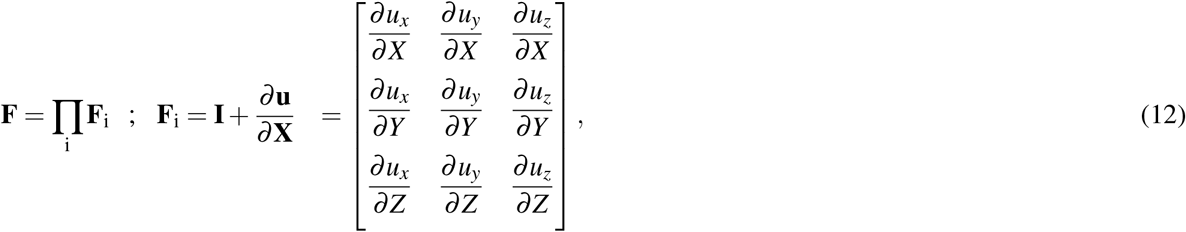

where u_x_, u_y_, and u_z_ are the displacements in the x,y, and z directions between two consecutive time frame, and the subscript *i* represents each image. The resulting Cartesian strains were converted to the widely used radial-circumferential-longitudinal (RCL) axes using an orthonormal transformation matrix (**Q**) as:

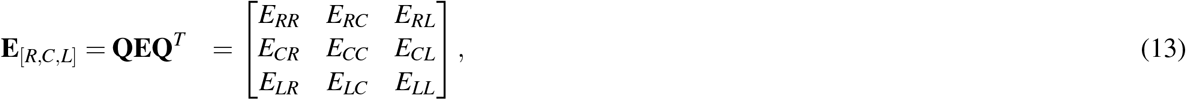

where *E*_*RR*_, *E*_*CC*_, and *E*_*LL*_ are the Lagrangian descriptions of radial, circumferential, and longitudinal strains, respectively. Similarly, the counterpart Eulerian strains are represented as *ε*_*RR*_, *ε*_*CC*_, and *ε*_*LL*_. The accuracy of NRIR in capturing strain patterns was validated against the ground truth (FE-derived strains) using mean squared error (MSE) analysis. Given the nonuniformity in data distribution between the phantom images and FE simulations, the unevenly distributed polar strains were mapped onto a target grid of fixed query points using radial basis function interpolation. For problem 2.1, the target grid for each slice was designed as a circular plane, and for problem 2.2, the grid was designed to mimic the American Heart Association (AHA) standard myocardial segmentation plot [42], with each target grid consisting of a two thousand query points. Thus, the MSE was calculated as follows:

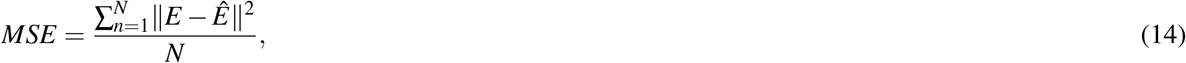

where *E* and *Ê* are the ground-truth and image-based strain quantities, respectively, and *N* is the total number of query points. MSE was evaluated at every increment for (i) three equidistant slices between *z* = 0 and *z* = 1 in the torsion problem 2.1, and (ii) the apical, mid, and basal slices in the LV problem 2.2. In addition, strain accuracy was evaluated using MSE for each segment in the AHA segmentation plot to evaluate the ability of image-derived methods to capture the regionality of cardiac motion.

### 3.6 Sensitivity analysis

A sensitivity analysis was conducted for the LV kinematics problem 2.2 to identify contributors to the deviations in strain accuracy. The accuracy of the NRIR algorithm in calculating strains was evaluated using the MSE metric for multiple configurations of the phantom. Key variables often known to contribute to inter-study and -operator variability in strain patterns and their effects on registration were considered. These variables included (i) through-plane resolution (slice thickness), (ii) in-plane resolution (grid spacing), (iii) temporal resolution (number of time frames), and (iv) noise intensity (Table. 2).

## 3 Results

### 3.1 Comparison of FE- and image-derived strains in the torsion problem

For the torsion problem (Section. 2.1), the FE-derived Eulerian strains were obtained at every node of the geometry and reported at the maximum twist angle (*θ*_*T*_ = 30°) over three sectional planes along its longitudinal axis (Figs. 4A and B). Regional variations in *ε*_*LL*_ and *ε*_*CL*_ were presented across all slices (Fig. 4A). The values decreased and increased incrementally from the inner to the outer radii of the cylinder at each slice, in the former and latter, respectively. Additionally, the strains at the inner and outer radii were equal to the analytically obtained strains for a thick-walled hollow cylinder under pure torsion (Eq. A1.6). Peak longitudinal contraction (*ε*_*LL*_ = − 0.13) and transverse circumferential-longitudinal shear (*ε*_*CL*_ = 0.26) took place along the outer radius of the cylinder.

**Fig. 4.**
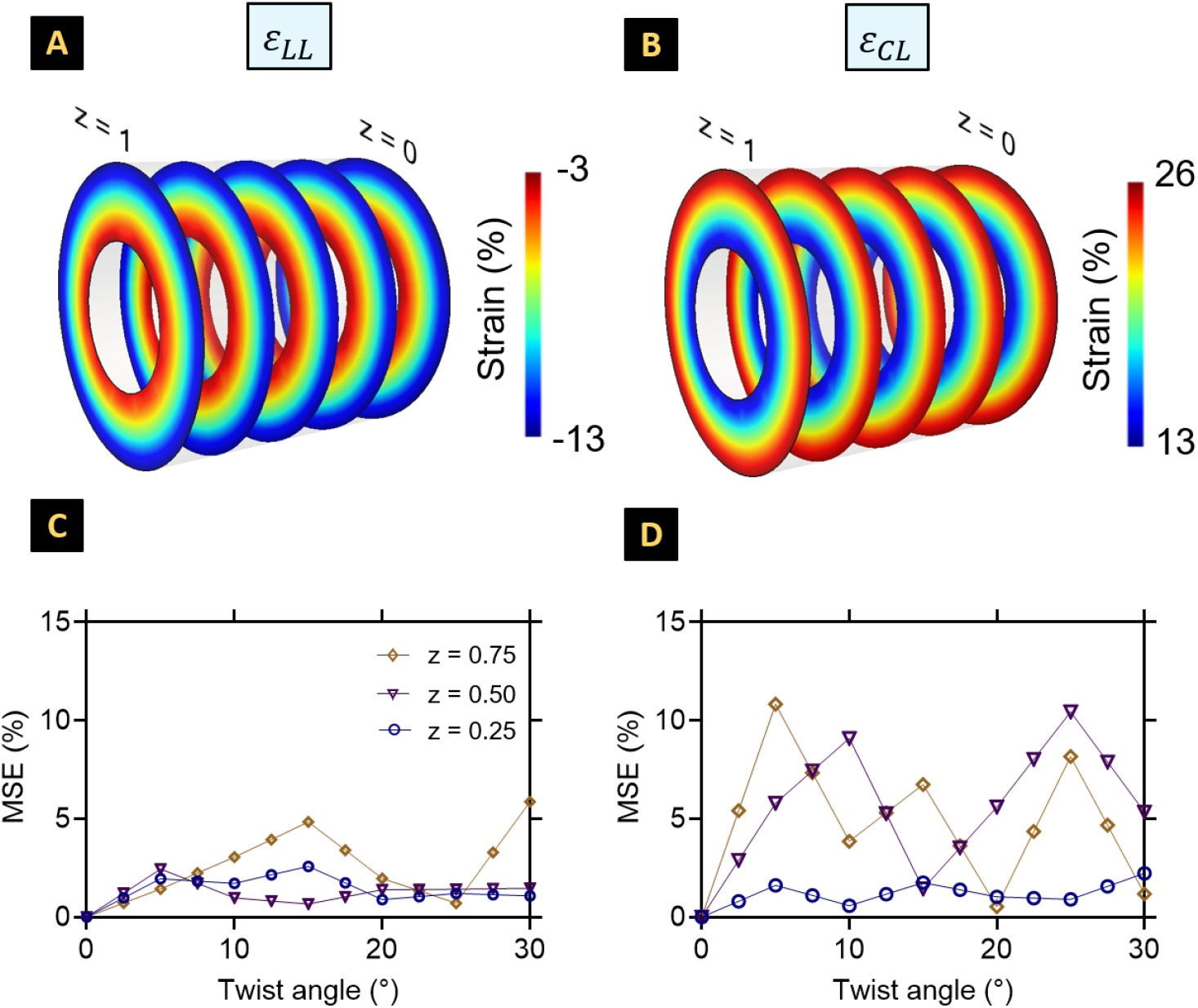
Finite element (FE) -derived strains along the (A) longitudinal axis and (B) the transverse circumferential-longitudinal plane for the thick-walled hollow cylinder under pure torsion at the maximum twist angle, *θ*_L_ = 60°. Strains are presented at three equidistant planes along the longitudinal axes. Mean squared error (MSE) curves calculated for FE- vs. image-derived (D) longitudinal, and (F) circumferential-longitudinal shear strains for the cylinder subjected to pure torsion at thirteen time frames between twist angles *θ*_T_ = 0° and *θ*_*T*_ = 30°. MSE readings are presented at three sectional planes along the geometry spaced 0.2 mm apart for the phantom configurations presented in Table 1. Each time frame corresponds to a single load increment, with the phantom consisting of 50 unique time frames.

The FE- (ground-truth) and image-derived strains were compared using MSE analysis at thirteen time frames between the untwisted and maximum twisted states of the cylinder. The strain errors are presented as a function of the twist angle at three equidistant sectional planes. Maximum errors for the normal strains (*ε*_*LL*_) were limited to 5% (Fig. 4C). Maximum deviations between the FE- and image-derived strains were observed in the *ε*_*CL*_ calculations, stemming from the middle and top slices at the early and later stages of twisting (Fig. 4D). The comparisons for *E*_*CL*_ revealed a maximum error of 10.45% at z = 0.75 mm, with all error values converging to under 5% at maximum twist (Fig. 4D).

### 3.2 Determination of “ground-truth” strains in the heart model

#### 3.2.1 Regional cardiac strain estimation using FE simulations

For the FE model of the murine LV (Section. 2.2), Lagrangian strains along the RCL axes were calculated at each node at ES relative to ED. The strain calculations are visualized across a LA plane passing through the center of mass of the LV (Figs. 5A-C) and three SA planes (Figs. 5D-F). The SA strains are presented as AHA segmentation plots across 16 segments of the LV base, mid, and apical slices. A strong presence of circumferential contraction and radial thickening was observed along the inferolateral LV, with similar contractile patterns in the septal LV also evidenced. These patterns were accompanied by marked regions of longitudinal contraction, and the strain values were consistent with the prescribed myofiber architecture, with the endocardium presenting greater contractility than the epicardium.

**Fig. 5.**
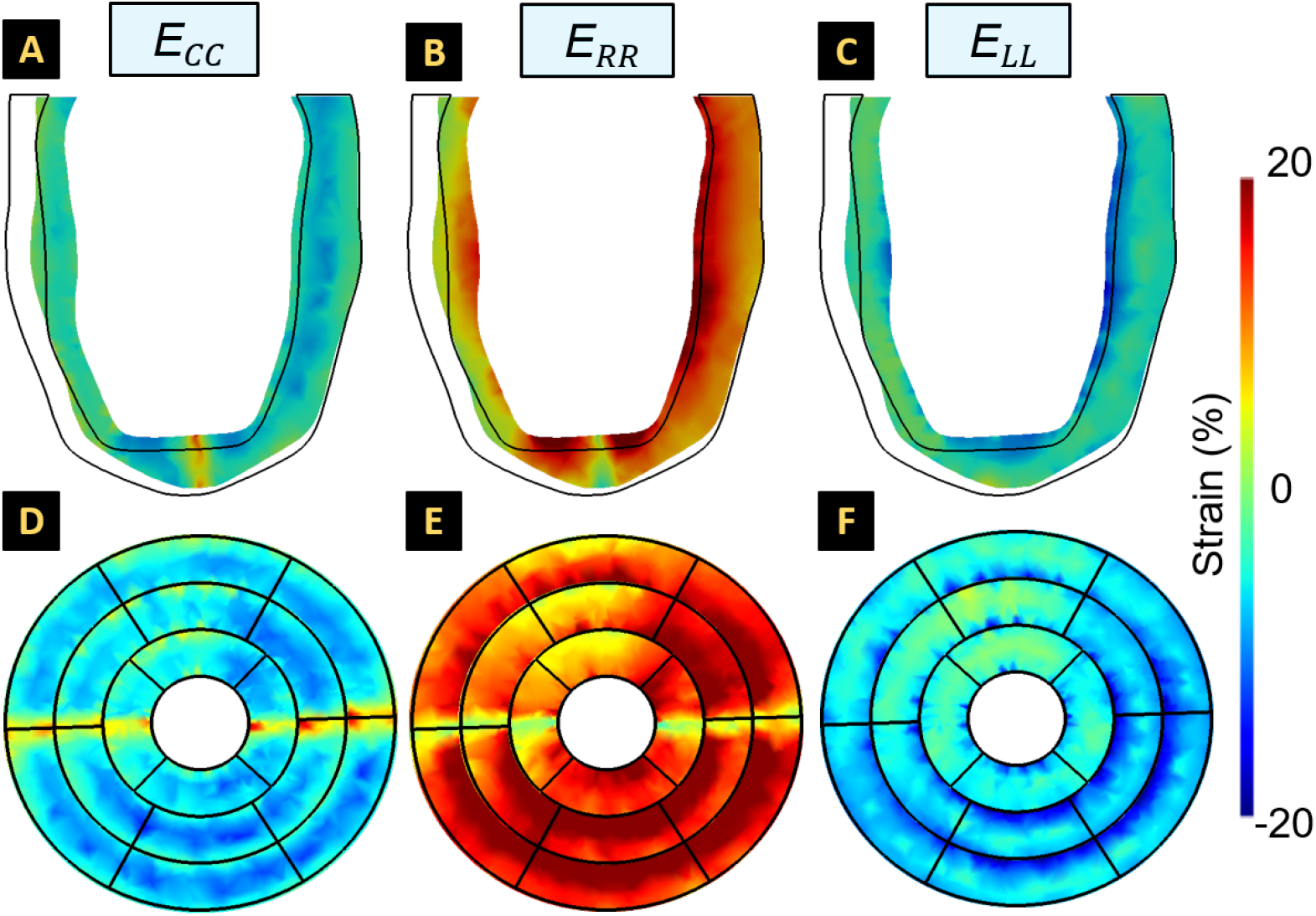
Finite element (FE) -derived (A-F) cardiac radial, circumferential, and longitudinal strains at end-systole (ES) relative to end-diastole (ED). The cardiac strains are presented along (A-C) an arbitrary long-axis plane and (D-F) short-axis (SA) basal, mid, and apical slices of the left ventricle. SA slices are presented as AHA segmentation plots, with the outermost and innermost rings corresponding to the basal and apical slices, respectively. The black outline in (C-E) denotes the LV geometry at ED.

#### 3.2.2 Comparison of cardiac strains with established motion benchmarks

A single global value of the benchmark strains was obtained by using the peak *E*_*CC*_, *E*_*RR*_, and *E*_*LL*_ values in each segment of the AHA segmentation plot. Peak LV contractility was noted at the inferior and lateral mid-apical slices (lateral: *E*_*CC*_ = −0.1847, and *E*_*LL*_ = −0.1752; septal: *E*_*CC*_ = −0.1058, and *E*_*LL*_ = −0.0811) and peak radial thickening was noted in the base and mid slices of the LV (*E*_*RR*_ =0.2148). These readings were consistent with existing literature on tissue-tagging-based calculations in healthy mice [43, 44, 45]. Liu et al. [43] confirmed peak contractility between the apex and mid slices of the LV, and among other studies, Chuang et al. [44] reported strain values of *E*_*CC*_ = −0.17±0.03 and *E*_*LL*_ = −0.17±0.04, respectively, using the DENSE algorithm. In addition to similar strain values, Zhong et al. [45] reported transmural variations in *E*_*CC*_ through the HARP algorithm, with maximum circumferential contractility observed in the endocardium of the LV. Similar observations were made in the contractile patterns reported by Zhong et al. and in this study, with septal strains estimated as *E*_*CC*_ = −0.08±0.03 and *E*_*RR*_ = 0.12±0.03. However, the literature on tagging-based calculations in mice is limited, the implications of which are expanded upon in the next section.

### 3.3 Comparisons between ground-truth and image-derived cardiac strains

MSE analysis was conducted between the FE- and image-derived cardiac strains at fifteen time frames between ED and ES. The errors are presented for all three strain quantities at the basal, mid, and apical slices of three phantoms with fixed in-plane spatial resolution and varying slice thickness values (Fig. 6). The temporal resolution of all phantoms was fixed at sixty total time frames to ensure that image-derived strains in each phantom were primarily influenced by the through-plane spatial resolution. The errors between FE- and image-derived *E*_*RR*_ and *E*_*CC*_ were mostly limited to 20% at ES for all phantoms (Figs. 6A-F). The maximum error increased with the anisotropy of the phantom (*w* = 1 vs. *w* = 4: *E*_*CC*_ = 17.563% vs. 23.265% and *E*_*RR*_ = 19.242% vs. 32.122%). In addition to the errors seen in the in-plane strain (i.e., *E*_*RR*_ and *E*_*CC*_) calculations, the thicker slices more significantly confounded motion tracking across the longitudinal axis, leading to the MSE as high as 30% observed at ES for all phantoms (Figs. 6G-I). To understand the association between image-derived strain calculations and through-plane resolution, SFR analysis was conducted (Supplementary Fig. S3). Whereas the isotropic phantom exhibited a moderately homogeneous spread of high and low spatial frequencies, the anisotropic phantom presented sharp attenuation in vertical frequencies. Such attenuation could potentially challenge the alignment of the fixed and moving images in the diffeomorphic demons algorithm (Eq. 9), thus creating errors in deriving through-plane strains (i.e., *E*_*LL*_). Nonetheless, despite deviations in the calculated strain values, the image-derived calculations captured the global contractile trend of the LV (Supplementary Fig. S4).

**Fig. 6.**
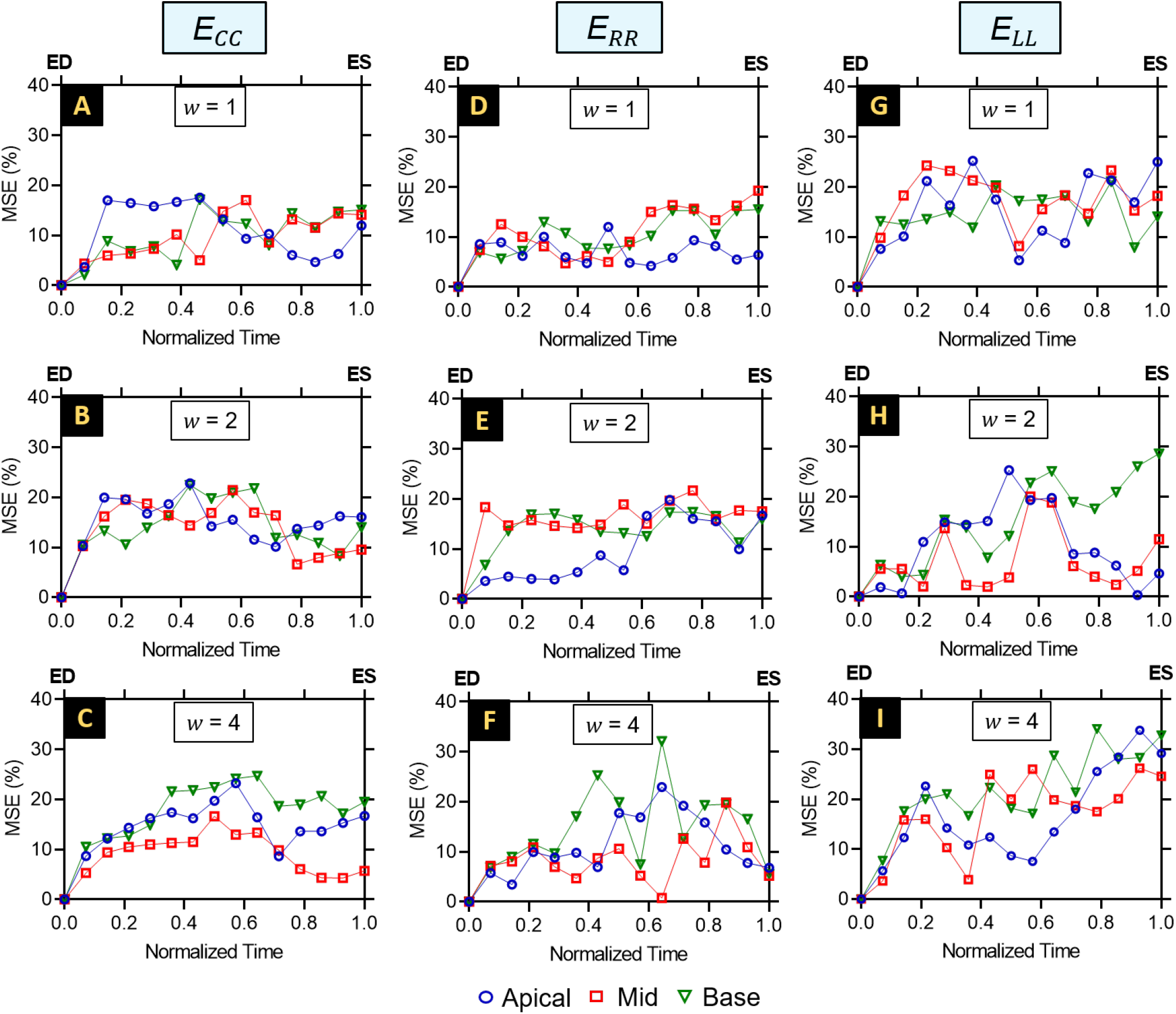
Mean squared error (MSE) curves calculated for FE- vs. image-derived regional cardiac strain quantities at 15 time frames (x-axis) between ED and ES. MSE readings are presented at the apical, mid, and basal slices of the in-silico phantom along the (A-C) circumferential, (D-F) radial, and (G-I) longitudinal axes. MSE curves are presented for phantoms of slice thickness, (top) *w* = 1 mm, (middle) *w* = 2 mm, and (bottom) *w* = 4 mm. Each time frame corresponds to a single load increment, with the phantom consisting of 60 unique time frames.

### 3.4 Effects of image quality on cardiac strain estimation

To further elucidate the effects of image resolution on strain calculations, MSE values are presented as error matrices for phantoms of various through-plane and temporal resolution parameters (Figs. 7 and 8). Considering the regions of peak contractility in the mid and apical slices, MSE analysis was more specifically examined in these regions (Fig. 7). Increasing the number of frames (i.e., improving the temporal resolution) reduced the MSE for all cardiac strain calculations irrespective of the anisotropy of the phantoms (Fig. 7). At the temporal resolution of 60 time frames, reducing the slice thickness improved cardiac strain calculations in both the mid and apical slices. While the temporal resolution was fixed at 15 time frames, in comparison to the *E*_*CC*_ calculations, the through-plane resolution of the phantom had a more pronounced influence over the estimation of *E*_*LL*_ in both the apical and mid slices of the LV (*w* = 1 vs. *w* = 4: *E*_*CC*_ = 31.20% vs. 25.00% and *E*_*LL*_ = 51.70% vs. 19.20%). Similar observations were made while analyzing the sensitivity of strain analysis to in-plane image features, such as in-plane pixel spacing and noise intensity (Fig, 8). While the reduction in pixel spacing improved the accuracy of the strain estimation, the addition of interpolation-driven noise due to resampling (Supplementary Fig. S2) or artifacts stemming from the through-plane reconstruction (Supplementary Fig. S5) may also contribute to additional errors in strain calculations.Thus, the MSE readings were influenced by factors such as blurring and resampling, underscoring the connection between enhancing image quality and reducing the spuriousness of image-derived strain calculations.

**Fig. 7.**
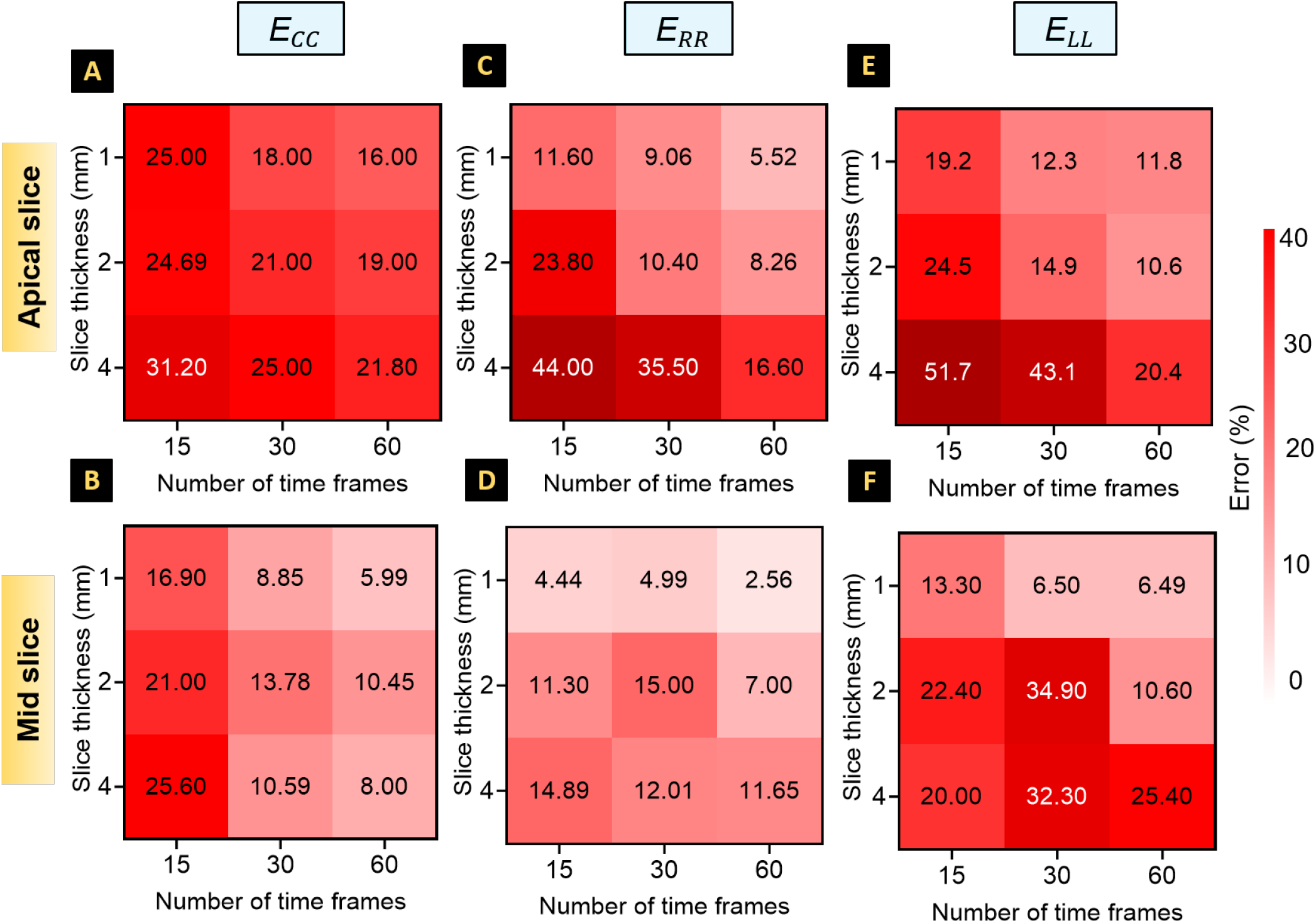
Error matrices describing the sensitivity of the regional strain calculations to slice thickness (through-plane resolution) and the number of time frames describing LV contraction from ED to ES (temporal resolution). Results are presented as their corresponding mean squared errors of image- vs. FE-derived strains at the (top) apical and (bottom) mid-slices of the phantom. The in-plane resolution of the phantom images was fixed at 1*×*1 *mm*^2^ with no noise.

**Fig. 8.**
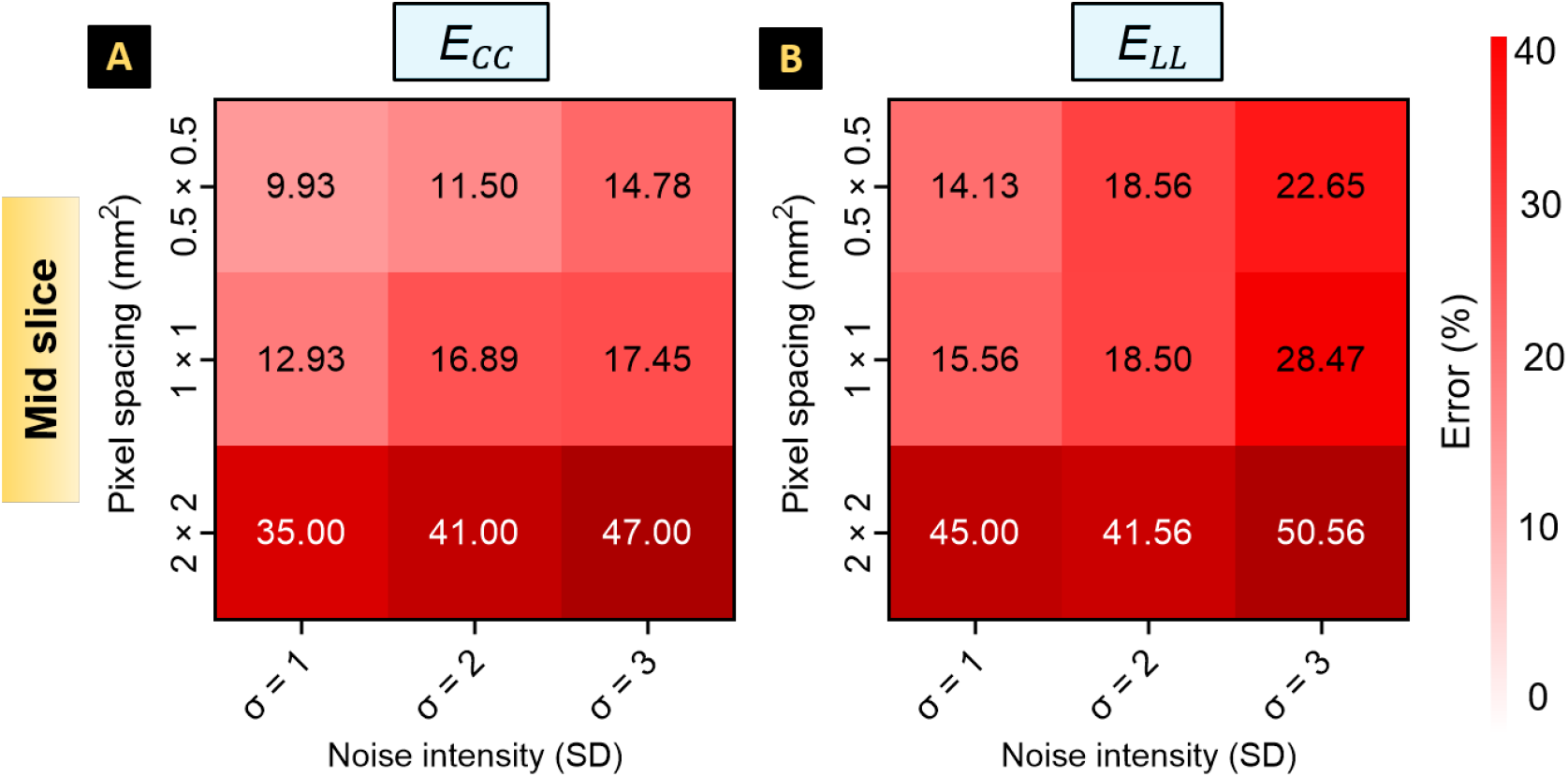
Error matrices describing the sensitivity of the regional strain calculations to pixel spacing (in-plane resolution) and additive noise. Results are presented as their corresponding mean squared errors of imaging- vs. FE-derived strains at the (A, B) mid-slice of the phantom. *σ*: standard deviation of the Gaussian kernel. The slice thickness of the phantom images was fixed at *w* = 1 *mm*

### 3.5 Transmural distribution of cardiac strain errors

While imaging parameters, such as resolution and noise, influence strain calculations, the regionality of cardiac motion arising from the complex organization of the LV microstructure is expected to impact strain estimation. The accuracy of the NRIR algorithm in assessing regional strain variations was analyzed by calculating the MSE over each segment of the AHA plot (Fig. 9). The analysis was conducted for all through-plane configurations of the phantom after fixing the temporal resolution to sixty total time frames. Upon examination, the concentration of spurious strains was identified in specific regions of the LV. Peak errors were exhibited in the mid and basal slices, coinciding with the regions of peak radial thickening. However, the spuriousness of strains was greater in the septal regions (Fig. 9) despite lower tissue contraction in the septal wall (Fig. 5).

**Fig. 9.**
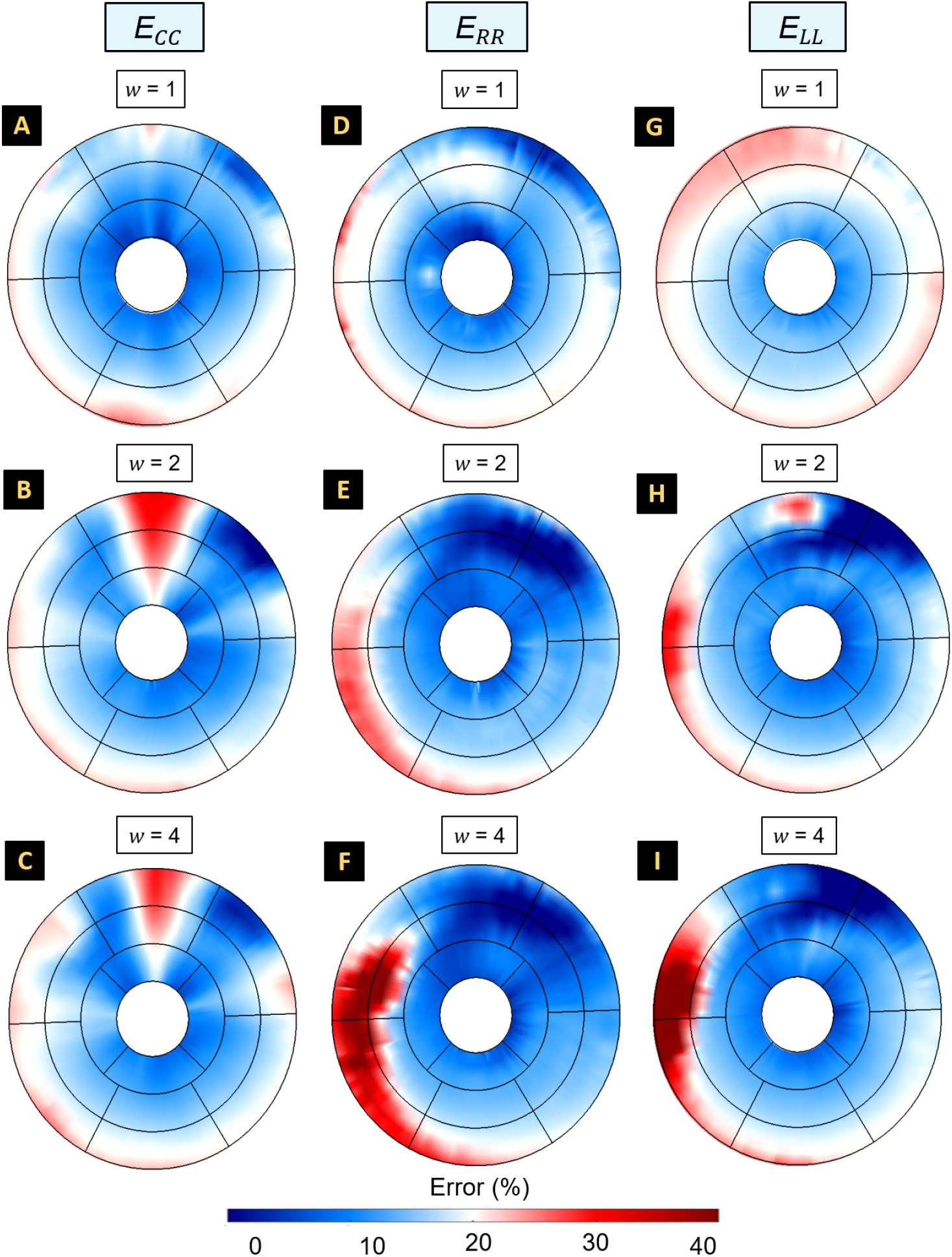
Heat maps illustrating the mean squared error between the FE- and image-derived strains at every section of the base, mid, and apical slices of the left ventricle at end-systole. MSE readings are presented at the apical, mid, and basal slices of the in-silico phantom along the (A-C) circumferential, (D-F) radial, and (G-I) longitudinal axes. MSE curves are presented for phantoms of slice thickness, (top) *w* = 1 mm, (middle) *w* = 2 mm, and (bottom) *w* = 4 mm. The temporal resolution of the phantom was fixed at 60 total time frames. Noise was added by varying the standard deviation (*σ*) of the normal distribution, and was limited to *σ* = 1.

## 4 Discussion

### 4.1 A comprehensive kinematic benchmark for cardiac strain validation

Despite the promising utility of motion-tracking algorithms in in-vivo strain estimation [11, 16, 46], the calculation of 4D regional stains remains limited. The reliability of regional strains as biomarkers, influenced by factors associated with image acquisition and motion calculations, hamper their full clinical translation. These limitations are particularly of concern in diagnosing regional injuries due to cardiac diseases. Due to the highly heterogeneous nature of cardiac motion (Fig. 5), strain calculation algorithms are susceptible to the generation of unrealistic strain quantities due to phenomena such as tag fading, motion artifacts, and poor resolution. These challenges are exacerbated in small animal models where the heart beats at a higher rate than in human patients. A few recent studies have demonstrated the potential benefits of in-vitro phantoms to validate CMR-based calculations of cardiac strains [24, 47] and stiffness [17]. However, in-vitro studies are challenged by the integration of the complex geometric and material properties of the heart, such as the myofiber organization, to replicate precise physiological ventricular motion while also having access to ground-truth kinematics. Our methodology then combined standard image processing techniques such as image registration, resampling, and blurring with statistics to streamline the validation of regional strain calculations against the “ground-truth” FE-derived strains. The benchmark was first tested using a simple thick-walled hollow cylinder under pure torsion to evaluate the validity of FE simulations in capturing the distinct torsional mechanics of the LV [48]. The regional variations in *ε*_*LL*_ and *ε*_*CL*_ distribution between the inner and outer radii of the cylinder challenged the accuracy of the image-derived Eulerian strains (Fig, 4). Subsequently, a similar methodology was implemented to create an in-silico phantom of the heart, with complex modes of motion used to test the accuracy in estimating 4D strain calculations using NRIR. Several errors were observed in the cardiac strain calculations, with image quality contributing to errors in *E*_*RR*_, *E*_*CC*_, and *E*_*LL*_ calculations.

### 4.2 Relationship between image quality and regional strain accuracy

The association of regularization terms with motion-tracking approaches has been known to be a significant source of variability. Subsequently, investigations were conducted to identify imaging parameters that influence the NRIR algorithm, and the findings were presented in terms of MSE and sensitivity analyses. Various phantoms were synthesized using a sinusoidal wave function to mimic standard cine CMR images with varying temporal and spatial resolutions. The diffeomorphic demons algorithm used herein applies Gaussian regularization to the displacement vectors to minimize the influence of sharpness and preserve the continuity of the solution [41]. Motion estimation was also influenced by geometric errors stemming from sharp image edges at the tissue-pool interface. For instance, sharp edges were produced as a consequence of through-plane reconstruction and in-plane interpolation (Supplementary Figs. S2 and S5). This substandard image resolution was expected to distort displacement derivatives, thereby producing errors in image registration. Thus, a strong correlation between edge sharpness and the generation of spurious strains was noted (Fig. 7). Accordingly, the consequential outcome of the global regularization was the underestimation of strain values, especially at the tissue-pool interface of the endocardium, and was confirmed by notable variations in the strain peaks (Supplementary Fig. S4) and the regional deformation patterns (Fig. 9). Thus, the heterogeneity in deformation over different segments of the same slice may challenge motion tracking via NRIR and thus contribute to errors.

Additionally, the slice thickness of the phantoms was observed as the primary contributor to errors in all strain quantities, with temporal resolution mitigating errors in *E*_*RR*_ and *E*_*CC*_. Interestingly, MSEs did not monotonically accumulate towards the ES, and higher MSEs occurred both within the ED-ES range and at ES, depending on the strain quantity, the SA slice position, and the phantom (Fig. 6). Despite improvements to the through-plane and temporal resolution, spurious strains were encountered in all phantoms. A potential contributor to strain errors could be the geometric misalignment of the LV between the SA slices due to the inherent curvature of the murine heart (Fig. 3). Despite the inherent errors introduced by the demons algorithms, the utilization of the kinematic benchmark with complex loading and geometries within the FE-derived phantoms presents valuable opportunities for the enhancement of imaging-based strain calculation.

### 4.3 In-silico phantoms offer enhanced recapitulation of in-vivo cardiac physiology

Muscle fibers in the myocardium are organized in helices, and fiber orientations within the helices are structured so that shortening of the fibers results in optimal ejection of blood and a fair distribution of load on the fibers.

Therefore, capturing the architecture and biomechanical behavior of native myocardium may serve as a crucial step in developing in-vitro cardiac models [26, 49]. However, materials used to fabricate cardiac phantoms, serving as in-vitro benchmarks for cardiac imaging studies, are often isotropic gels, making the incorporation of intricate native organization of the myofiber network a challenging task. Furthermore, in-vitro phantoms are challenged by the infeasible incorporation of P-V loops, usually loaded via simple rotational and tensile forces. These limitations hinder investigations into the regionality of cardiac motion under complex kinematic forces such as twisting, shearing, and contracting. Indeed, the physiological motion of the heart is characterized by key modes of motion during different phases of the cardiac cycle, such as in-plane biaxial stretching during diastole and wall thickening and torsion during systole. In this study, we presented a completely in-silico CMR phantom incorporating subject-specific geometry, fiber architecture, and P-V measurements to recapitulate the various modes of motion.

We addressed the limitations of traditional phantom-based validation approaches by employing a well-established transversely isotropic computational model of the LV. The material model (Eq. 2), in conjunction with mouse-specific fiber architecture and P-V loop boundary conditions, generated regionally varying contractile patterns (Figs. 5D-I), which enabled the investigations of the effects of cardiac strain regionality on NRIR. For instance, distinct regional variations in tissue strains were between the inferolateral regions of the LV and the anteroseptal wall (Fig. 5). As such, sources of error were observed in both the septal and lateral regions, with error potentially stemming from the significantly varying tissue strains between different sections of the LV (Fig. 9). Thus, by varying the material, architectural, and geometric parameters in the FE simulations, the specificity of the in-silico phantom to in-vivo physiology in health and disease. For example, the material constants in Eq. 3 can be altered to describe isolated infarction zones within the LV [50]. Such investigations can provide insight into the sensitivity of motion-estimation algorithms to non-contractile zones (as a result of scar formation) and provide higher fidelity in regional strain calculations. Additionally, the model can be easily extended to provide benchmark motion in humans through similar methods described within this study (Appendix. A2).

### 4.4 Implications and future direction

The innate complexity of regional cardiac motion, resulting from intricate myocardial architecture and passive and active phases of motion [26], remains a prominent challenge in rigorously quantifying full 4D cardiac motion in vivo. Thus, the ability to mitigate strain errors is paramount while evaluating the performance of any imaging-based calculation. As presented in this study, errors originate from a combination of geometric and motion effects on image quality. The complex nature of 4D ventricular kinematics demands the establishment of a quantification and validation methodology. However, the rigorous quantification of 4D myocardial strains remains a challenging problem and an area of promising research. Potential improvements in estimating regional strains can be obtained by enhancing the spatial discretization of the ventricular geometry through high-resolution imaging or by conducting super-resolution reconstruction using standard cine images. Other alternatives include implementing physics-informed regularization terms [51] and machine learning methods [52] that bridge the gap between the parametric and non-parametric transformation terms in motion tracking algorithms. The methodology presented herein facilitates a feasible manipulation of material, geometrical, and architectural parameters to synthesize an extensive dataset of images. We expect to create a vast dataset in training for tasks such as classification, segmentation, and motion estimation to provide a comprehensive physics-informed approach to estimating 4D cardiac motion. Furthermore, we expect the in-silico heart phantom to provide the foundation to directly characterize soft tissue parameters through standard image-based motion analysis. Through the in-silico heart phantoms presented in this study, steps were taken toward validating cardiac strain calculations in health and disease. By manipulating the material and geometric parameters of the phantom, conditions such as hypertrophy and infarction can be easily accommodated in the in-silico phantom, thus providing potential improvements in the clinical translation of 4D cardiac strains. The harmonious integration of image processing and FE simulations can provide a comprehensive approach comprising subject-specific mechanical characteristics to enhance the fidelity of regional strains as biomarkers and assist in clinical intervention.

### 4.5 Limitations

Despite the high potential clinical utility of the in-silico cardiac phantom, a few limitations exist in the study. First, our study was limited to two computational models of LV kinematics. While this study intended to introduce a new perspective into designing kinematically intricate phantom models, further investigations that incorporate additional subject-specific geometrical, architectural, and mechanical parameters obtained from diffusion tensor imaging, histology, and P-V measurements can further deconvolute the various contributors to spurious cardiac strain estimation. Moreover, the in-silico heart phantom can also be extended to the other facets of cardiac function, such as the right ventricle [53, 54], wherein ground-truth strains in health and disease are yet to be established [55], thus creating a tailored kinematic benchmark for the entire heart. Second, image artifacts were restricted to SNR modeled as a normal distribution during the image formation process, with limited control over regional maps. Operator-dependent motion artifacts arising from the inherent complex interactions between blood flow and the cardiac wall can influence image quality. Thus, we expect region-specific texture maps that provide a more realistic description of the various tissue elements to facilitate a more versatile benchmark for motion estimation algorithms. We anticipate texture generation methods that implement mathematical models of scattering (ultrasound) and gradient distribution based on tissue excitation (CMR) can provide even more realistic synthetic images. Despite the absence of artifacts beyond the Gaussian noise, the in-silico phantoms facilitated investigations that could differentiate between the various image-related contributors to error, thus offering an excellent benchmark to assess and improve the reliability of existing cardiac strain estimation algorithms.

## 5 Conclusion

In this study, we presented a novel in-silico heart phantom that facilitates a comprehensive validation of regional cardiac strains estimated from medical imaging data. To the best of our knowledge, this is the first application of a completely in-silico FE-derived phantom to enhance the characterization of imaging-derived regional cardiac strains. The phantom enhances the fidelity of current standards in validation protocols by offering physiological cardiac kinematics together with ground-truth values for validation. We provided a methodology to facilitate the generation of phantoms of various configurations that can be easily implemented in investigating the fidelity of motion estimation algorithms in evaluating different key modes of motion in the cardiac cycle. Our investigations focused on mouse-specific geometry and loading to test the accuracy of NRIR via diffeomorphic demons to estimate regional strains during cardiac contraction (i.e., ED to ES). Various contributors to spurious strain estimation were investigated, with potential sources of error validated against a comprehensive kinematic benchmark. Our findings revealed significant deviations between the benchmark cardiac strain calculations and strains derived from the phantom images. Potential contributors to these deviations were investigated, with both image quality and strain regionality concluded to have challenged the accuracy of cardiac strain estimation. Ultimately, a detailed representation of 4D cardiac kinematics was investigated, with such analysis offering the potential to include high-fidelity 4D regional contractile patterns of the LV in standard clinical screening of cardiac function. Our future objectives are to create an extensive dataset to facilitate the inclusion of the in-silico phantom to improve the standard of image-based motion analysis and provide a comprehensive physics-informed approach to estimating 4D cardiac motion.

## 6 Acknowledgements

R. Avazmohammadi was supported by the NHLBI grant R00HL138288. J. Ohayon was partially supported by the Agence Nationale de la Recherche (ANR), through the SIMR project (ANR-19-CE45-0020). T. Mukherjee was supported by the American Heart Association (AHA) predoctoral fellowship 24PRE1240097.

## 8 Data availability

The data that support the findings of this study are available from the corresponding author, R.A., upon request.

## Appendix

### A1 Analytical solution of a hollow cylinder under pure tosion

Since the exact solution of a thick-walled hollow cylinder under pure torsion is dependent on the material model, geometry, and boundary conditions, a kinematic representation of a cylinder subjected to torsional shear is summarized below. Let us consider an arbitrary hollow cylinder of radii, a ≤ R ≤ b, and height 0 ≤ Z ≤l in the undeformed polar space. Here, R and Z correspond to the radial and longitudinal axes in the undeformed state, with Θ corresponding to the circumferential axis. The plane Z = l is rotated about the Z axis, while Z = 0 remains fixed. The combined deformation of the geometry can be represented as:

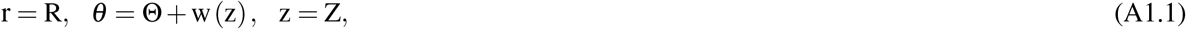

here, r, *θ*, and z correspond to the polar coordinates in the deformed state, and w(z) represents the rotational twist about the z axis. Since there is no extension, i.e., z = Z, the rotation can be conceptualized as a twist at various plane sections of the cylinder. Considering the applied twist per unit length, *θ*_T_ = 0 at z = 0 due to the base remaining fixed, the deformation can be rewritten as,

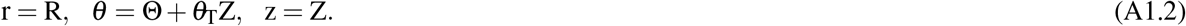

The deformation gradient, **F** can be derived using this representation as:

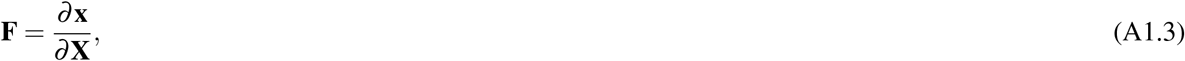

here **x** and **X** are the coordinate systems in the deformed and undeformed states, respectively. In this case, **x** = (*r, θ, z*) and **X** = (*R*, Θ, *Z*) are represented in the polar coordinates. Thus, **F** is derived as:

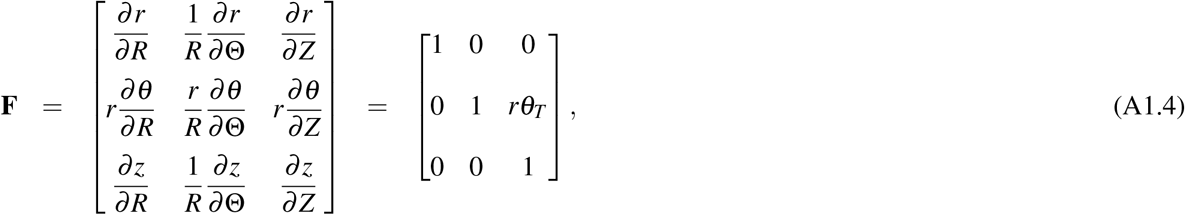

the accompanying Eulerian Cauchy-Green strain tensor, *ε* (Eq. 10) can be derived using **F** and the identity tensor **I** as:

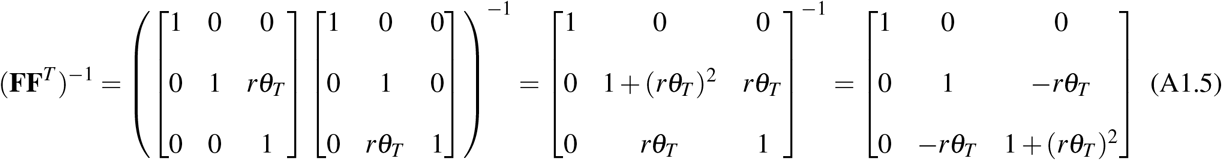

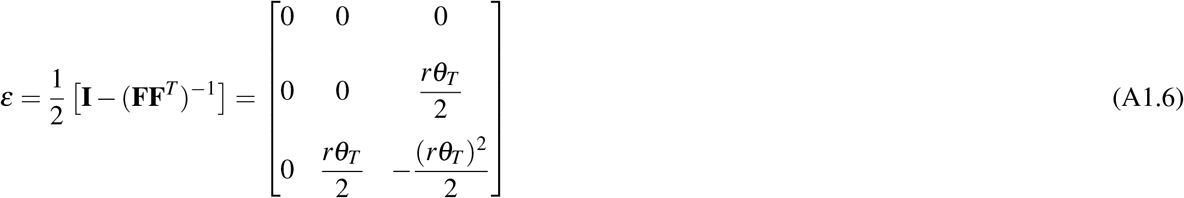

### A2 Human-specific in-silico heart phantom

#### A2.1 Forward simulations

The phantom synthesis approach was implemented to create a human-specific in-silico phantom to highlight the proposed utility of the method in the clinical translation of 4D regional strain calculations. The human-specific phantom was developed using CMR images obtained for a human patient presenting left ventricular diastolic dysfunction. Eight SA scans were captured using a 3.0-T clinical scanner (Siemens Verio; Siemens, Erlangen, Germany) with phased-array coil systems. Segmentation and reconstruction were performed at the end of isovolumic relaxation, i.e., early diastole (EaD), using the steps described in section 2.2.1. The 3D LV geometry was truncated below the valve plane and meshed using linear tetrahedral elements (Fig. A1A). In the absence of patient-specific fiber architecture, synthetic myofibers were applied to the model, with the preferred fiber angle varying transmurally from − 60° at the epicardium to 60° at the endocardium

The LV geometry was modeled as passive myocardium using a hyperelastic, transversely anisotropic, and nearly incompressible material taking the form of a modified exponential Fung model (*W*^*dev*^ in Eq. 3). The material constants, *B*_1_, *B*_2_, and *B*_3_ in Eq. 4 were taken from a previous study and the stiffness parameter in Eq. 3 was determined as *c* = 296.0 Pa. Finally, the LV model was loaded to an end-diastolic pressure *EDP* = 9.0 mmHg (Fig. A1A). The resulting circumferential and radial strains (*E*_*CC*_ and *E*_*RR*_) at ED relative to EaD, were taken as the “ground-truth” values (Fig. A1B). A single global measure representing the strains was obtained by taking the mean of all strains in the basal, mid, and apical slices of the LV.

#### A2.2 Phantom synthesis and strain comparison

Synthetic phantom images of the human-specific LV model were created using the steps described in section 2.3. Briefly, the displacement vectors describing the deformation of the LV from EaD to ED were used to create individual phantoms for every load increment. Subsequently, the nodes in 3D space were rasterized onto 2D images. Images of the following parameters were created at eight sections of the LV: spatial resolution of 200*×* 200 and temporal resolution of 1 frame/increment. No artificial noise was added to the images (Fig. A1C). NRIR was conducted, and strains were calculated using the methods described in sections 2.4 and 2.5, and comparisons were made between the global strains obtained from the FE model and the synthetic phantom images (Fig. A1).

**Fig. A1.**
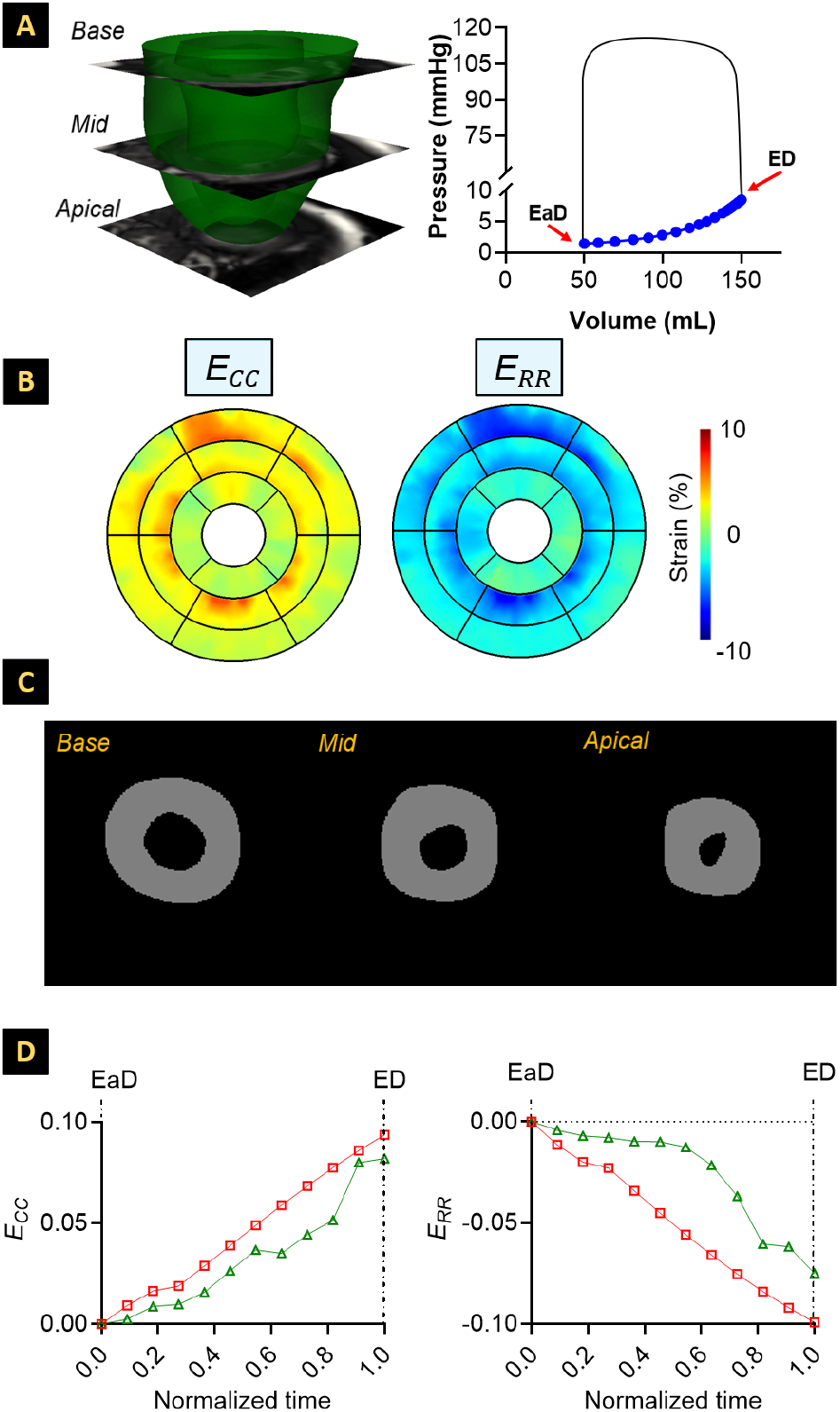
(A) 3D human-specific finite-element (FE) model for the passive relaxation of the left ventricle (LV). The endocardium of the LV geometry (left) was loaded using the filling pressures described in the pressure-volume curve for an end-diastolic pressure of 9 mmHg (right). (B) FE-derived cardiac circumferential and radial strains at end-diastole (ED) relative to early-diastole (EaD). (C) Grayscale images of the in-silico phantom at various short-axis sectional planes of the LV. (D) Comparison of the global strain estimation at various short-axis sectional planes of the LV at various stages of deformation between EaD and ED along the circumferential (left) and radial axes (right). The global strains are estimated as the peak individual strain in the LV.

## Supplementary material

**Supplementary Figure S1.**
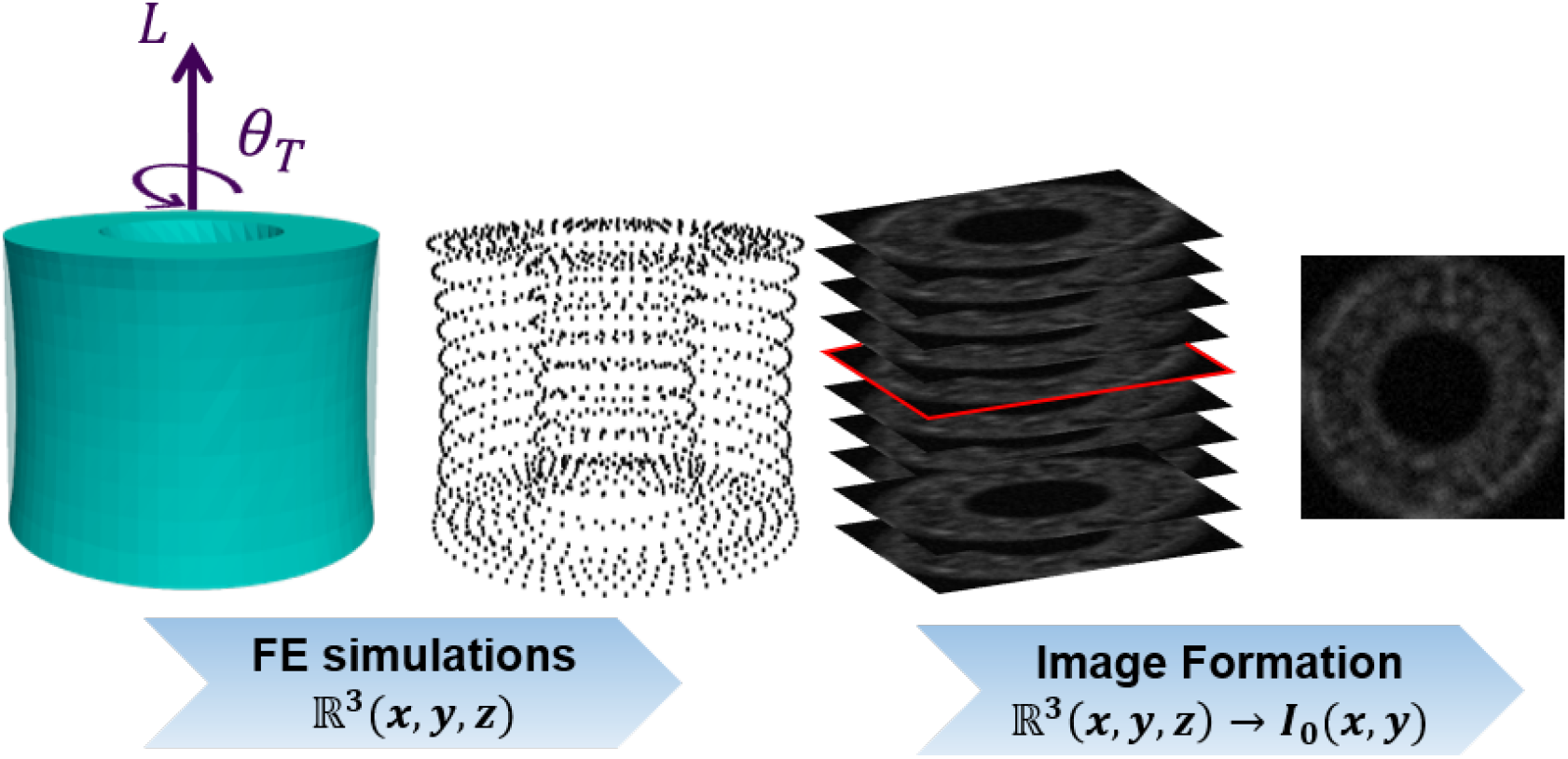
Schematic outlining the stages in creating synthetic phantom images from finite element simulations of a cylinder under pure torsion. The images comprise the following parameters: spatial resolution = 1 *×* 1 mm^2^, slice thickness (*w*) = 1 mm with no added noise. Max torque angle, *θ*_T_ = 60°

**Supplementary Figure S2.**
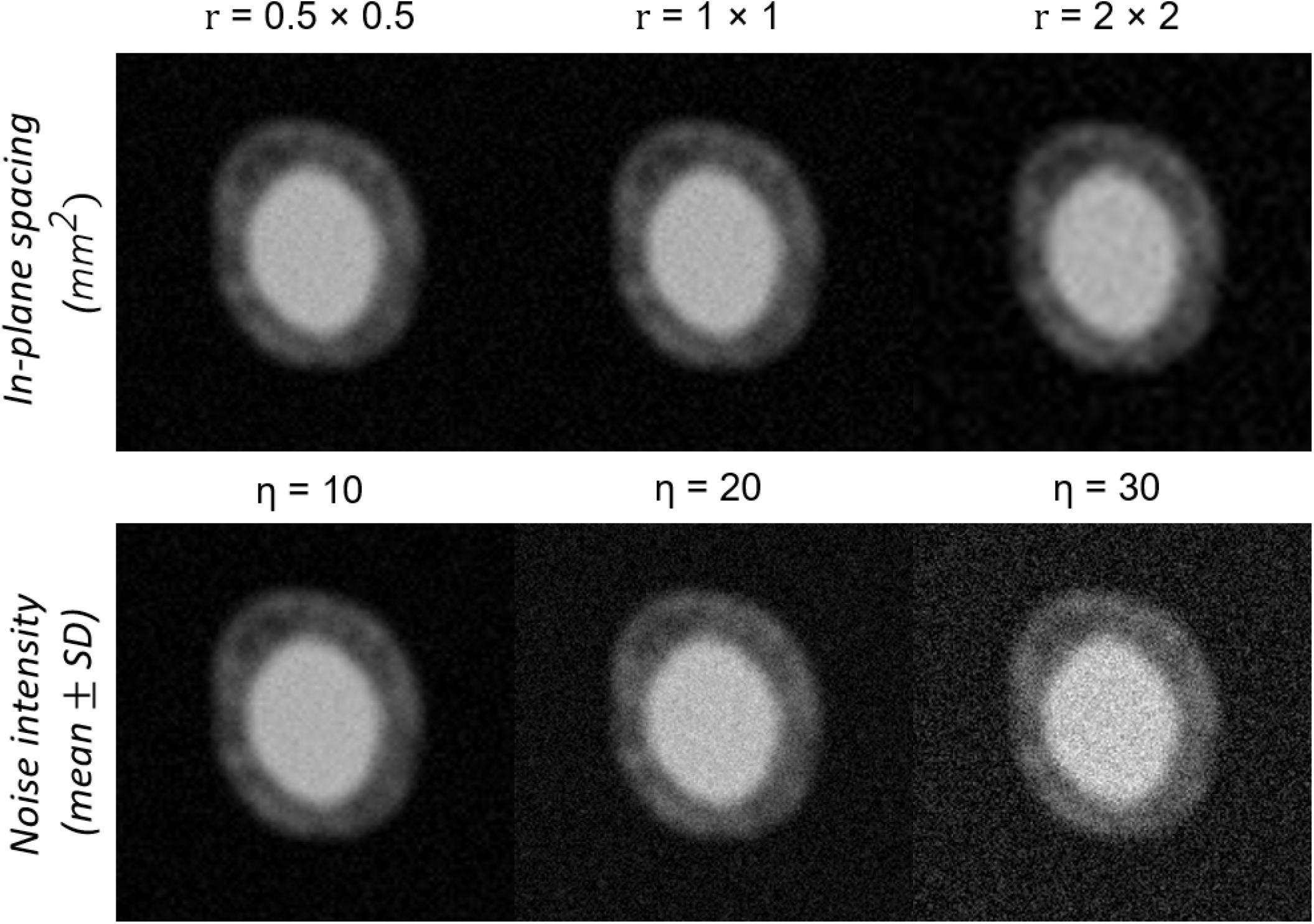
Grayscale images at an arbitrary short-axis slice of the left ventricle demonstrating the effects of adding noise via (top) resampling and (bottom) normal distribution of the mean gray level intensity. These images correspond to a phantom of the following parameters: spatial resolution = 1 *×*1 mm^2^, slice thickness (*w*) = 1 mm, and temporal resolution = 1 frame/load increment.

**Supplementary Figure S3.**
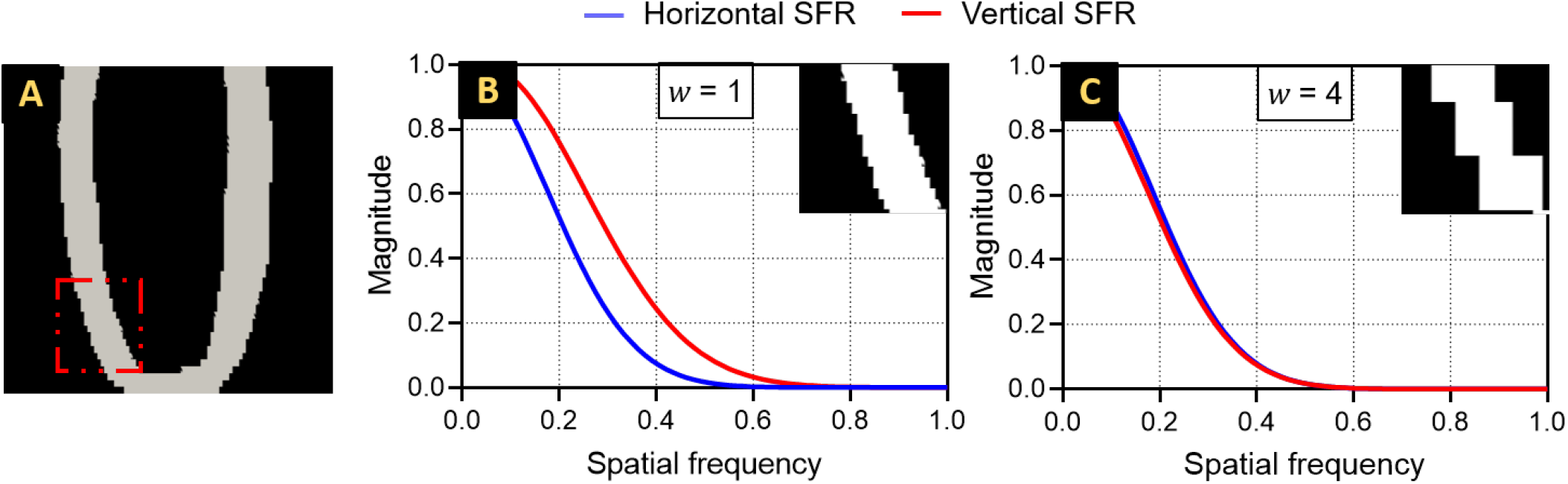
Spatial frequency response (SFR) for an arbitrary long-axis plane sampled for two distinct configurations of the cardiac phantom. (A) Interrogation windows of 150 *×*150 pixels are used to measure the spatial frequencies at the mid-apical section of the septal wall. The SFR curves are presented for phantoms with slice thickness, (B) *w* = 1 mm, (C) *w* = 4 mm. The overlap between the SFR curves indicates equal sharpness of the image in the horizontal and vertical directions of the image. Horizontal and vertical distributions of the SFR are shown in blue and red, respectively.

**Supplementary Figure S4.**
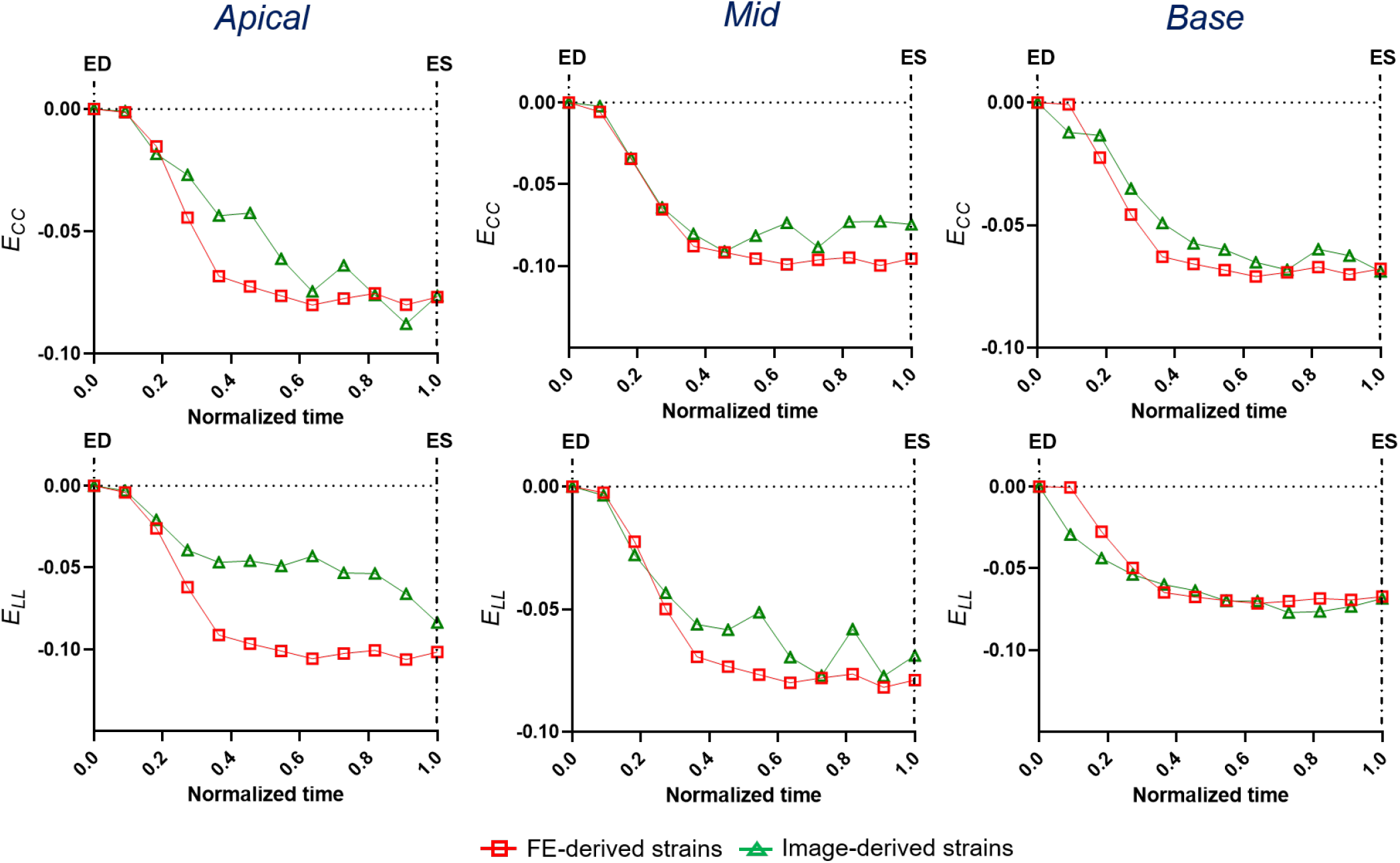
Comparison of the global strain estimation at various short-axis sectional planes of the left ventricle at various stages of deformation between end-diastole (ED) and end-systole (ES) along the (top) circumferential and (bottom) longitudinal axes. The global strains are estimated as the mean of all individual strains at each LV slice. The NRIR-derived strains are represented for a phantom of the following parameters: spatial resolution = 1*×* 1 mm^2^, slice thickness, *w* = 1 mm, and temporal resolution = 1 frame/load increment. *E*_*CC*_: circumferential and *E*_*LL*_ longitudinal strains.

**Supplementary Figure S5.**
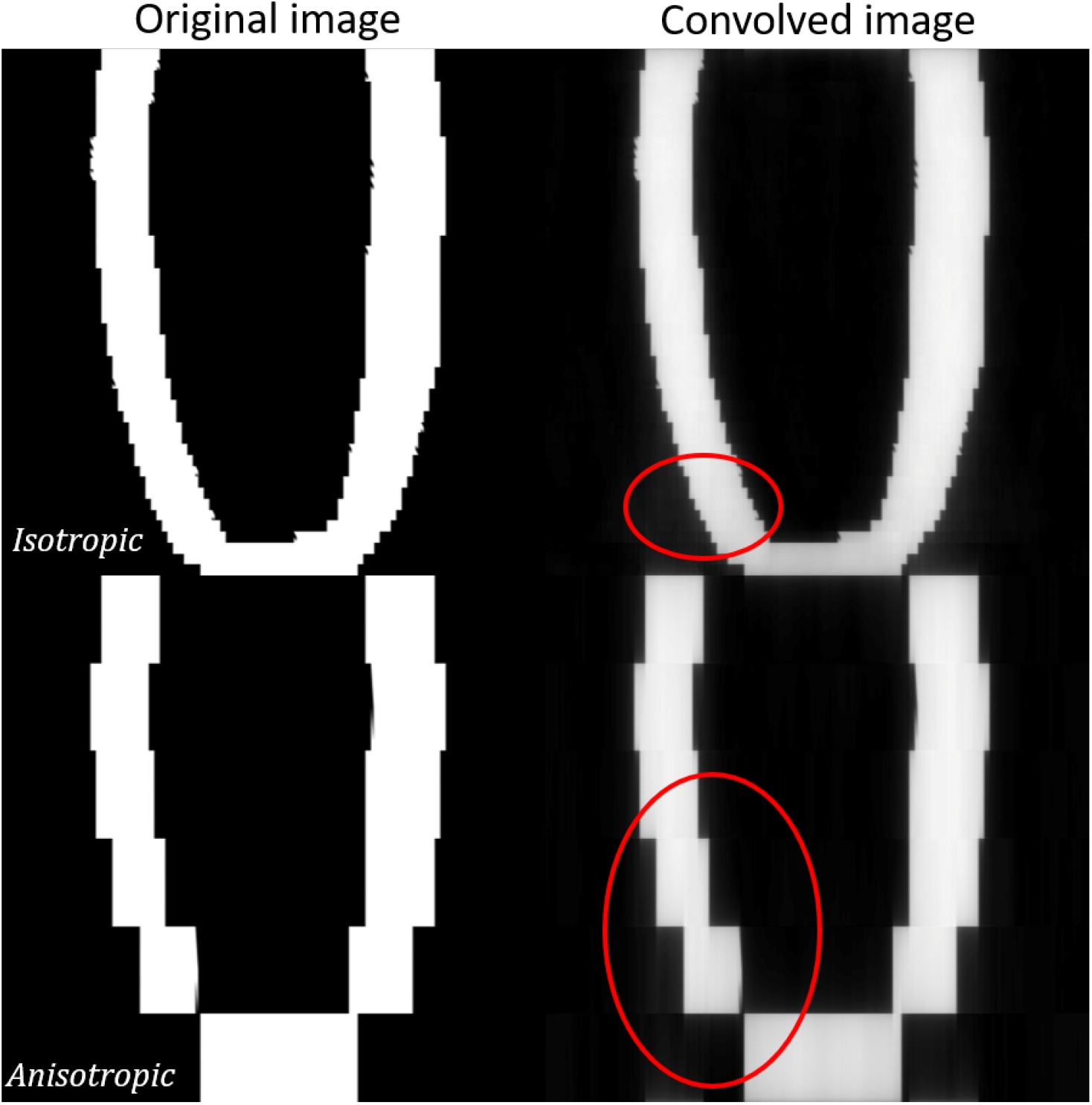
Representative images of the through-plane long-axis (LA) reconstruction of the LV by stacking short-axis slices leading to the generation of grayscale artifacts. (Left) the original LA reconstruction, (right) convolution of the original image with its corresponding Fourier transform. The regions circled in red denote areas influenced by the sharp image edges and are analogous to standard motion artifacts. *w*: slice thickness

